# GNEval: A Context-Specific Benchmark for Gene Network Inference with Single-Cell Foundation Models

**DOI:** 10.64898/2026.11.25.769274

**Authors:** Tianxing Hu, Weiwei Wu, Zhaohui S. Qin, Chang Su

## Abstract

Gene network (GN) inference plays a critical role in understanding gene relationships and their roles in biological functions and diseases. Existing single-cell GN inference methods often rely on limited data from individual studies. Single-cell foundation models, pretrained on large and diverse datasets, provide a range of gene-gene association signals that offer a promising basis for GN inference. However, it remains unclear which GN properties these signals capture and how they compare with domain-specific methods. To address this gap, we introduce **GNEval**, a unified benchmark for evaluating pretrained model derived gene associations for network inference. We benchmark 21 gene representations and association scores from 11 models, including static gene embeddings (S), contextual embeddings (C), and gene-gene attention scores (A). GNEval assesses inferred networks based on their consistency with functional and perturbational evidence, including 14 Perturb-seq datasets across diverse cell systems, perturbation types and data sizes. The results reveal substantial heterogeneity across evaluation tasks and models. Rankings based on Gene Ontology and Perturb-seq evidence were nearly uncorrelated (Spearman *ρ* = −0.05). Context-independent representations showed a substantial advantage in recovering functional relationships (ΔAUROC = 0.13), whereas context-dependent gene associations improved recovery of perturbation-associated differential expression relationships by 3.8×. The domain-specific method achieves relatively strong performance on both evidence types, offering a better balance across the two evaluations than most foundation-model-derived networks. Overall, gene representations and association scores show task-specific strengths, highlighting the need for task-aware selection and integration with domain-specific methods.

## 1 Introduction

Gene network (GN) inference emerged with the advent of high-throughput transcriptomic profiling (Stuart et al., 2003) and have since become an indispensable tool for understanding how genes coordinate to perform biological functions and how dysregulation of biological pathways may underlie disease. For example, weighted gene co-expression network analysis (Langfelder and Horvath, 2008) has been widely used across tissues and disease conditions, with successes in neuro-science (Geschwind and Konopka,2009; Gandal et al.,2018), cardiometabolic disease (Chen et al., 2008), and cancer (Segal et al., 2004). In the single-cell era, methods have also emerged to infer GNs from single-cell RNA-seq (scRNA-seq) datasets, revealing how genes coordinate within specific cellular contexts (e.g., Su et al. 2023; , Kamimoto et al. 2023), . Throughout this paper, we use GN to refer to undirected gene association networks, commonly known as co-expression networks, in which edges represent statistical association between genes rather than direct or directed regulatory relationships; we discuss this scope and its limitations in Section 4.

However, a common limitation of existing single-cell GN inference methods is that they rely solely on the dataset at hand, which is often limited in sample size. Consequently, these approaches may not fully leverage the extensive prior knowledge of gene relationships accumulated across single-cell data atlas (Regev et al., 2017; CZI Cell Science Program et al., 2025) and scientific literature.

Recent advances in single-cell foundation models (scFMs) (Theodoris et al., 2023; Cui et al., 2024; Yang et al., 2024; Hao et al., 2024; Kalfon et al., 2025; Hu et al., 2026) and large language models (LLMs) (Chen and Zou 2025), provide a new opportunity to advance GN inference for small-scale, context-specific datasets. For example, pretrained static gene embeddings may encode functional similarities between genes, contextual gene representations may capture cell-state-dependent gene relationships, and attention scores may reflect model-internal gene-gene associations. Specifically, these models encode rich prior knowledge from large-scale single-cell datasets and literature-curated resources in the learned gene representations, and similarity of gene representations inform similarity of gene function (Hu and Wei, 2026). We therefore hypothesize that scFMs can be highly valuable for learning gene networks in small-sample settings, such as on a new dataset collected by a clinical collaborator for a specific question of interest, and that they can complement domain-specific methods.

To evaluate this hypothesis, it is crucial to understand what GN information scFMs encode. Three questions are open: what relationships do scFMs capture, where are they captured (static embeddings, contextual embeddings, or attention scores), and how do they compare with classical domain-specific statistical and computational methods applied to the target data alone? These questions are hard to answer because gene relationships are context-dependent and heterogeneous, and because different evaluation strategies capture distinct types of network information.

Existing benchmarks do not resolve these questions. Benchmarks of scFMs on gene-related tasks (Wenteler et al., 2025; Wu et al., 2025; Zhong et al., 2025; Li et al., 2025; Radig et al., 2026; Hou et al., 2026) rarely include GN inference and focus largely on context-independent representations; comparisons against domain-specific GN inference methods are scarcer still. Benchmarks of GN inference (Pratapa et al., 2020; Chevalley et al., 2025; Karamveer et al., 2026; Nourisa et al., 2026), in turn, do not systematically assess scFM-derived signals across biological contexts and typically rely on a single source of evidence. To our knowledge, no benchmark has systematically evaluated context-specific gene embeddings or attention-derived association scores from scFMs for GN inference against perturbational evidence from the matching cellular context.

Advances in genome editing and scRNA-seq, particularly Perturb-seq across diverse cellular contexts and perturbation types, provide new opportunities to study gene relationships at scale (Dixit et al., 2016; Norman et al., 2019; Tian et al. 2021;, Schmidt et al., 2022). Perturb-seq measures cellular transcriptomic profiles upon perturbation of regulators, such that genes differentially expressed between perturbed and control cells are potential targets of the perturbed regulator in GNs. Furthermore, Perturb-seq captures both perturbation-induced gene effects and matched unperturbed transcriptomic profiles from the same cellular context, allowing scFMs to be applied to the unperturbed cells and their inferred GNs to be evaluated against experimentally measured perturbational effects.

To bridge these gaps, we introduce a comprehensive benchmarking framework **GNEval** for evaluating gene representations and association scores in GN inference (Figure 1). GNEval covers 14 datasets, 11 models, and three types of gene representations and association scores, and evaluates gene relationships using complementary evidence from gene function database and experimental perturbational evidence. GNEval assesses network quality from three perspectives: edge-set enrichment, edge ranking, and recovery of supported gene relationships. We employ unified data preprocessing, standardized representation extraction, consistent evaluation metrics, and sensitivity analyses to ensure fair and comprehensive comparisons. We additionally examine stability across benchmark settings, generalizability across biological contexts, and computational cost and its scaling with cell count.

**Figure 1:**
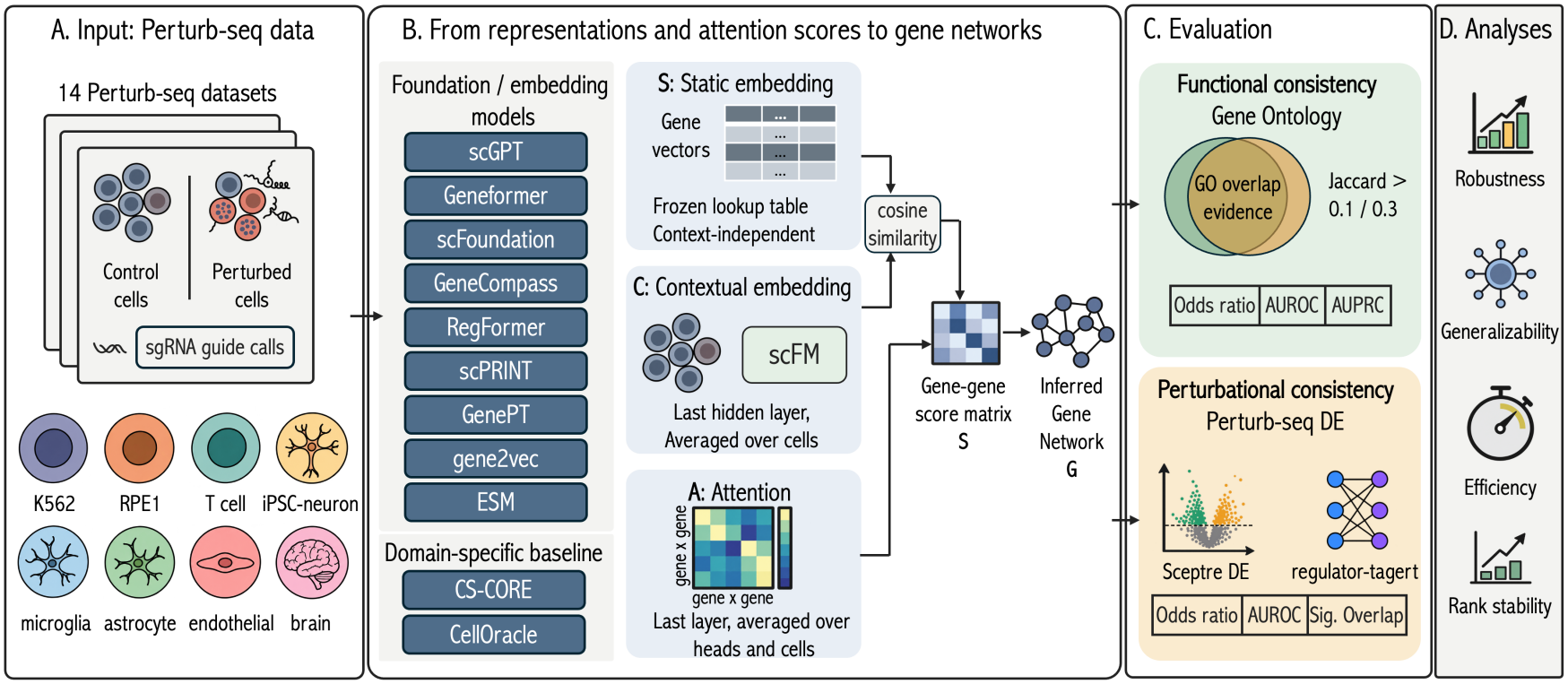
Overview of GNEval.

Through systematic experiments, we derive several key insights. First, context-independent and context-dependent representations capture distinct aspects of gene relationships. Context-independent representations generally better recover functional relationships defined by GO, whereas context-dependent representations and association scores better recover perturbational evidence. Second, the domain-specific method achieves relatively strong performance on both evidence types, offering a better balance across the two evaluations than most scFM-derived networks.

Our main contributions are as follows. First, to our knowledge, GNEval is the first systematic benchmark of context-specific gene associations derived from scFMs for GN inference. We evaluate diverse scFMs and other pretrained gene representation models against domain-specific GN inference methods across biological contexts. Second, to our knowledge, GNEval introduces the first systematic use of Perturb-seq data for context-matched evaluation of GNs, combining perturbational and functional evidence across a curated collection of 14 latest Perturb-seq datasets with analyses of robustness, generalizability, and computational efficiency. Moreover, we develop a standardized pipeline that harmonizes heterogeneous Perturb-seq datasets and derives perturbation-based evidence for gene relationships. Finally, we provide an extensible and reproducible implementation for benchmarking GNs, supporting future evaluation and development of gene representation and network inference methods. (https://github.com/tianxinghu/GNEval-benchmark)

## 2 GNEval Framework

### 2.1 Task Definition: Gene Network Inference

Let *V* = {*g*_1_,. .., *g*_*N*_} denote a set of genes. We propose to infer GN using a gene-gene association matrix

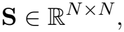

where *S*_*ij*_ quantifies the inferred association between genes *g*_*i*_ and *g*_*j*_.

**S** can be derived from different model outputs. For gene embeddings, each gene *g*_*i*_ is associated with an embedding **z**_*i*_ ∈ ℝ^*d*^, and pairwise association scores are derived from the learned representations, using cosine similarity (Yang et al., 2024; Chen and Zou, 2025; , Hu et al. 2026),

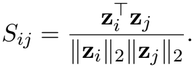

For pairwise gene-gene scores obtained directly from models, including attention scores and association scores from domain-specific methods, we use their inferred scores as **S** (Section 2.2).

To enable comparison across methods, gene pairs are ranked according to their inferred association scores, and the top *K* ranked pairs are retained to form the inferred network

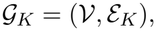

where *V* denotes the set of evaluated genes and *E*_*K*_ contains the *K* highest-ranked gene pairs according to their absolute association scores. We consider *K* ∈ {5,000, 10,000, 20,000}.

The benchmark evaluates whether the resulting networks recover biologically supported genegene relationships using independent functional and perturbational evidence.

### 2.2 Models

GNEval evaluates GNs inferred from 11 models spanning diverse sources (Table 1). We include four recent scFMs, **scGPT** (Cui et al., 2024), **Geneformer** (Theodoris et al., 2023), **scFoundation** (Hao et al., 2024), and **GeneCompass** (Yang et al., 2024), which were trained on atlas-scale single-cell data. We extract signals at different stages of the model, including context-independent pretrained token embeddings, contextualized final-layer gene embeddings, and attention scores. To provide broader baselines, we further consider gene representations learned from alternative data sources beyond single-cell transcriptomics. **GenePT** (Chen and Zou, 2025) provides literature-derived gene embeddings generated using OpenAI’s text-embedding-ada-002 model from the summary sections of the NCBI Gene database, while **gene2vec** (Du et al., 2019) learns gene representations from co-expression pairs derived from microarray-based gene expression data using a Word2Vec objective. We also include **ESM** (Lin et al., 2023), a protein language model from which gene-level token embeddings are constructed by averaging the pretrained representations of all proteins encoded by each gene, following the construction adopted in scFMs (Rosen et al., 2026; Kalfon et al., 2025). Beyond general-purpose representation models, we evaluate other two recent scFMs, **RegFormer** (Hu et al., 2026), and **scPRINT** (Kalfon et al., 2025), with architectures and training strategies tailored toward modeling gene regulatory relationships.

**Table 1:** Methods under comparison in GNEval. **Used as: S** static gene embedding, **C** contextual embedding, and **A** gene-gene attention read directly as edge scores.

| Method | Dim | Type | Training basis | Used as |
|---|---|---|---|---|
| GenePT | 1536 | Text embedding | GPT text of gene descriptions | S |
| gene2vec | 200 | Word2Vec | Bulk co-expression (GEO) | S |
| ESM | 5120 | Protein LM | ESM2 embeddings of gene products | S |
| scGPT | 512 | Transformer | scRNA-seq (∼33M cells) | S, C, A |
| scFoundation | 768 | Transformer | scRNA-seq (∼50M cells) | S, C, A |
| Geneformer | 768 | Transformer | scRNA-seq (∼95M cells, gc-104M) | S, C, A |
| GeneCompass | 768 | Transformer | scRNA-seq (∼120M human + mouse cells) + prior knowledge | S, C, A |
| RegFormer | 512 | Mamba (state-space) | scRNA-seq (∼26M human cells), GRN-ordered | S, C |
| scPRINT | 256 | Transformer | scRNA-seq (∼50M cells) | C, A |
| CS-CORE | — | Co-expression estimator | scRNA-seq (no pre-training) | Domain-specific |
| CellOracle | — | GRN inference | scRNA-seq + motif-based GRN prior (no pre-training) | Domain-specific |

Finally, we include two domain-specific methods: **CS-CORE** (Su et al., 2023), which infers gene-gene relationships directly from observational scRNA-seq, with its test statistic used as the association score, and **CellOracle** (Kamimoto et al., 2023), which reconstructs context-specific gene regulatory networks (GRNs) by combining scRNA-seq with motif-based transcription factor (TF)-target priors and regularized regression, with the regression coefficient used as the association score.

### 2.3 Gene Association Scores and Network Construction

GNEval evaluates three broad classes of methods (Table 1): scFMs, other pretrained gene representation models, and domain-specific GN inference methods. For each scFM, we extract available gene representations or gene-gene association scores from different model components and categorize them into three types: static embeddings (**S**), contextual embeddings (**C**), and attention scores (**A**) (Baek et al., 2025). Because individual scFMs can provide multiple representation types, the 11 evaluated models yield a total of 21 representations or association scores. In contrast, other pretrained models based on non-single-cell data, such as GenePT, gene2vec, and ESM, provide only static embeddings and do not produce dataset-specific contextualized representations. Finally, CS-CORE and CellOracle directly infer gene-gene associations from single-cell data without pretraining and are therefore included as domain-specific baselines. Table 1 summarizes the representation or score types available from each evaluated model.

#### Static embeddings (S)

Static embeddings are context-independent gene representations obtained without providing any dataset-specific single-cell observations to the model. Each gene is therefore associated with a single fixed vector for a given model, shared across all downstream datasets. For scFMs, these representations are extracted from the pretrained token embedding table when available. We additionally consider context-independent representations derived from other sources, including GenePT (Chen and Zou, 2025) embeddings generated from gene-description text, gene2vec (Du et al., 2019) embeddings learned from bulk gene co-expression, and ESM (Lin et al., 2023) protein embeddings aggregated to the gene level. All these representations do not depend on the single-cell datasets used for downstream evaluation. These embeddings are converted to pairwise gene-gene association scores using cosine similarity.

#### Contextual embeddings (C)

Contextual embeddings are dataset-specific gene representations obtained by running each scFM in inference mode on single-cell expression profiles. For each dataset, we extract gene representations from the final hidden layer and average the representations of each gene across cells in which that gene is observed. These embeddings are converted to pairwise gene-gene association scores using cosine similarity and treated as context-dependent.

#### Attention scores (A)

For models with self-attention, we directly extract gene-gene attention scores by running each scFM in inference mode on single-cell expression profiles. We use the attention matrix from the final layer, average across attention heads, and then average each gene pair over cells in which both genes are observed. These scores are used directly as association scores and treated as context-dependent.

To extract contextual embeddings and attention scores from scFMs, we use all non-targeting control cells of each dataset (capped at 12,000 for the two largest screens (Zhu et al., 2026; Shi et al., 2026), sampled proportionally across donors and lanes), the same cells CS-CORE is fit on. CellOracle is fit on all control cells. The pretrained parameters remain fixed, and the expression data are used only to contextualize the representations, without fine-tuning the model.

For the domain-specific methods, we use the CS-CORE test statistic and the CellOracle absolute regression coefficient as gene-gene edge scores. We construct networks by retaining the top 5,000, 10,000, and 20,000 highest-scoring edges for each method when available. Because CellOracle produces sparse networks restricted to TF-target pairs, it is evaluated at the network sizes it reaches together with its full network (Appendix E).

### 2.4 Functional and Perturbational Evaluation

GNEval evaluates inferred GNs against two complementary sources of biological evidence: GO-based functional relationships and Perturb-seq-derived perturbational relationships. We use GO annotations (Gene Ontology Consortium, 2004) to define co-functional gene pairs among the top 1,000 highly variable genes in each dataset. Treating GO as a bipartite gene-term graph, each gene *i* is represented by its annotated GO-term set *G*_*i*_, and pairwise functional similarity between genes *i* and *j* is quantified using the Jaccard index *J* (*i, j*) = |*G*_*i*_ ∩ *G*_*j*_|*/*|*G*_*i*_ ∪ *G*_*j*_| (Roohani et al., 2023). Gene pairs with *J* (*i, j*) *≥* 0.1 or 0.3 are considered GO-supported, with both thresholds evaluated separately to assess robustness.

For perturbational evaluation, Perturb-seq datasets were processed using a unified pipeline (see details in Appendix B). Differential expression (DE) analysis was performed with sceptre (Barry et al., 2021, 2024) using raw counts from single-guide cells, with non-targeting-guide cells serving as controls. Only the top 1,000 highly variable genes were considered in DE testing, with sequencing depth and detected-gene count included as covariates. An edge *i → j* is considered DE-supported when perturbation of regulator *i* produces a significant expression change in gene *j* with FDR < 0.05 and | log_2_ FC| > 0.25.

For both evidence sources, we use three metrics to capture complementary aspects of network quality. Two metrics assess the identification and ranking of biologically supported edges. For identification, enrichment is quantified using the odds ratio computed from the contingency table comparing inferred edges with supported gene pairs. This metric measures whether supported edges are overrepresented among the inferred edges. For ranking, AUROC is used to assess whether supported gene pairs tend to receive higher absolute association scores than other pairs. The third metric is evidence-specific: Because GO-supported pairs are typically sparse relative to the full set of gene pairs, GO-based evaluation additionally includes AUPRC to assess positive-pair recovery under class imbalance. For Perturb-seq-based evaluation, we further use the significant-overlap rate, defined as #{*r* ∈ *R*_test_ : *Q*_Fisher_(*r*) < 0.05}*/*|*R*_test_|, where *R*_test_ is the set of regulators retained after filtering (Appendix A) and *Q*_Fisher_(*r*) is the FDR-adjusted *p*-value from a one-sided Fisher’s exact test assessing over-representation of perturbation-derived DE genes among the inferred target genes of regulator *r*. This measures the agreement between inferred target genes and perturbation-derived DE genes across regulators.

### 2.5 Comparison of Representation Types

We tested whether the use of target-dataset context was associated with systematic differences in functional and perturbational consistency. Pretrained representations and association scores were grouped into context-independent and context-dependent categories, with CS-CORE and CellOr-acle analyzed separately as a domain-specific observational baseline (Table 1; Section 2.3). For each dataset, metric values were averaged within each category and compared across datasets using paired two-sided Wilcoxon signed-rank tests. Multiple-testing correction and additional statistical details are provided in Appendix H.

Additional analyses evaluated robustness to network size, GO similarity thresholds, cell sub-sampling, cell population, dataset characteristics, and computational scaling. (Appendices J.1 and K).

## 3 Results

### 3.1 GNEval Perturb-seq Datasets

We evaluate all GNs on 14 publicly available Perturb-seq datasets spanning diverse cell types, perturbation modalities, and dataset scales. To our knowledge, this is the first systematic curation of Perturb-seq datasets for benchmarking GN inference across biological contexts. We obtain the raw count matrices and associated metadata from the Gene Expression Omnibus (GEO) and develop a consistent preprocessing pipeline across datasets (see Section 4 for code availability). Detailed dataset characteristics, including the numbers of cells, perturbations, cell types, and significant DE edges are summarized in Table 2.

**Table 2:**
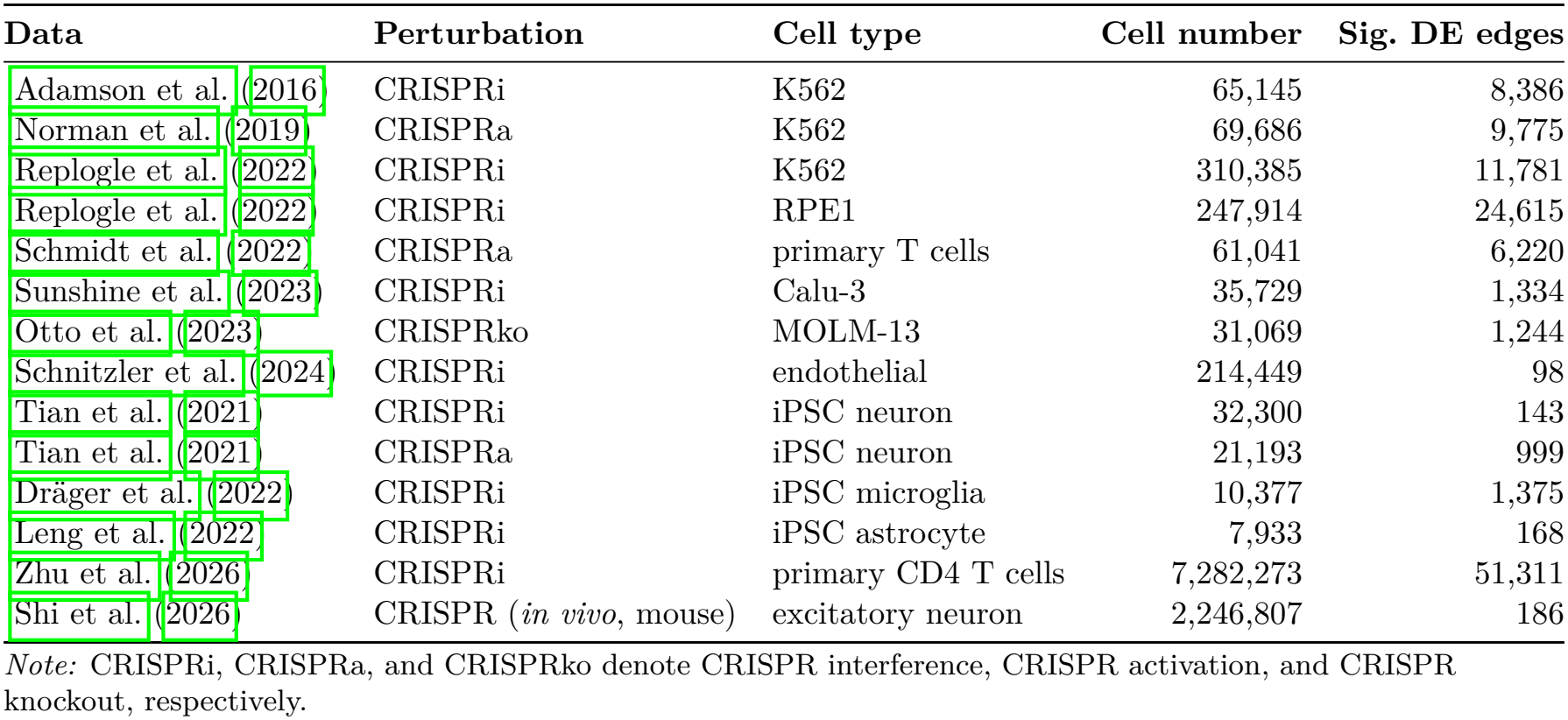
Perturb-seq datasets used for GNEval. “Sig. DE edges” is the number of significant regulator-gene pairs identified by sceptre (FDR < 0.05 and | log_2_ FC| > 0.25).

### 3.2 Functional and Perturbational Evaluation Performance

We evaluate gene association scores derived from scFMs and other pretrained gene representation models, together with CS-CORE and CellOracle, across 14 Perturb-seq datasets using complementary functional and perturbational evidence. GO-based evaluation measures consistency with functionally related gene pairs defined from GO annotations, with performance summarized in Figure 2. Perturb-seq-based evaluation measures consistency with regulator-target relationships supported by DE following CRISPR perturbation, with results summarized in Figure 3.

**Figure 2:**
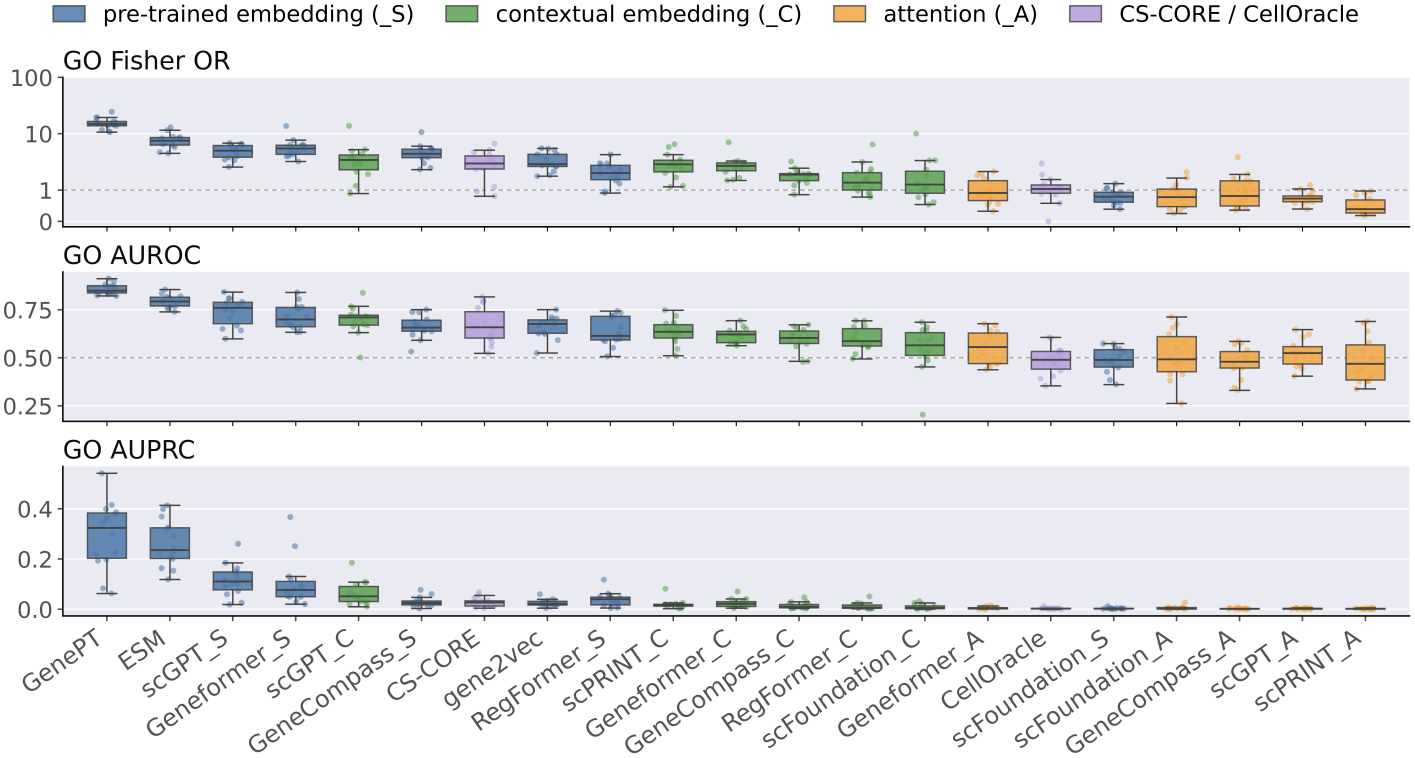
GO-based evaluation across 14 Perturb-seq datasets, measured by Fisher OR, AUROC, and AUPRC. Dashed lines indicate expectation under random chance. Methods are ordered by the average rank of their median performance across the three metrics.

**Figure 3:**
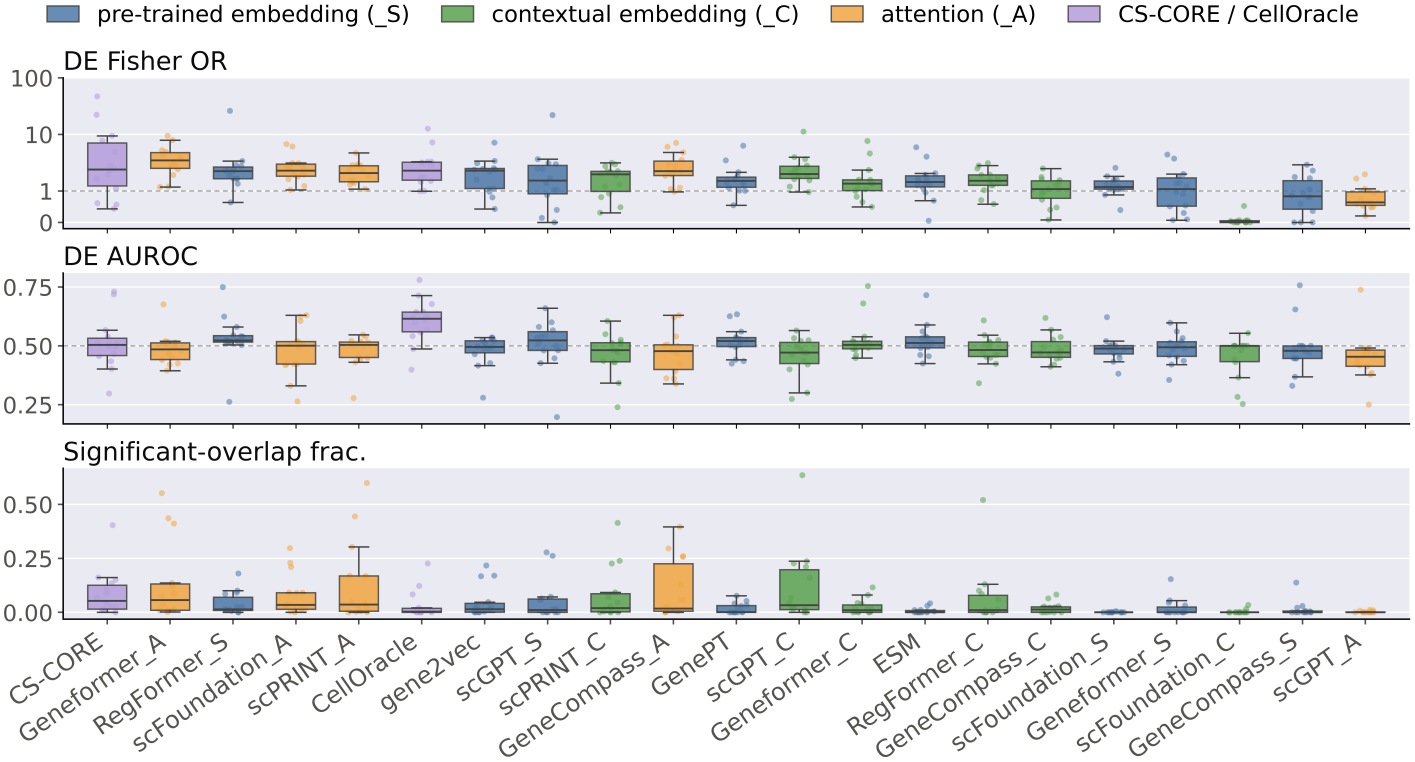
Perturb-seq-based evaluation across 14 Perturb-seq datasets, measured by Fisher OR, AUROC, and Significant Overlap Rate. Dashed lines indicate expectation under random chance. Methods are ordered by the average rank of their median performance across the three metrics.

In Figure 2, a clear overall trend is evident. GenePT leads every GO metric (median OR 14.9, AUROC 0.85, AUPRC 0.32), followed by ESM (7.5, 0.79, 0.24). Next come the static scFM embeddings scGPT_S and Geneformer_S. All attention scores, CellOracle and scFoundation_S sit at chance. In Figure 3, performance against Perturb-seq DE evidence is weak overall. By mean rank over the three metrics, the leaders are CS-CORE, Geneformer_A, RegFormer_S, scFoundation_A and scPRINT_A. Geneformer_A and CS-CORE have the highest DE odds ratios (median 3.5 and 2.4) and the largest fraction of regulators with significant overlap (median 5.7% and 5.3%). Median DE AUROC falls between 0.45 and 0.53 for every representation except CellOracle, the only method clearly above chance (median *≈* 0.62).

Method rankings are essentially uncorrelated between functional and perturbational evaluations (Spearman *ρ* = −0.05, *n* = 21, Figure 4). For each evaluation setting, methods were ranked separately on each of the three metrics based on their median performance across datasets, and the three metric-specific ranks were then averaged to obtain the overall rank for that setting. In Figure 4, the methods group by how the representation is built. Static gene embeddings (_S) occupy the GO-favored side: GenePT, ESM and scGPT_S are the top three on GO but rank 8th-14th on Perturb-seq. Attention scores (_A) and CellOracle occupy the opposite corner: Geneformer_A ranks second on perturb-seq, yet all _A scores rank 15th-21st on GO. Contextual embeddings (_C) fall in between. The near-zero correlation suggests that GO-based and perturbation-based evaluations capture different aspects of a GN, so neither can substitute for the other.

**Figure 4:**
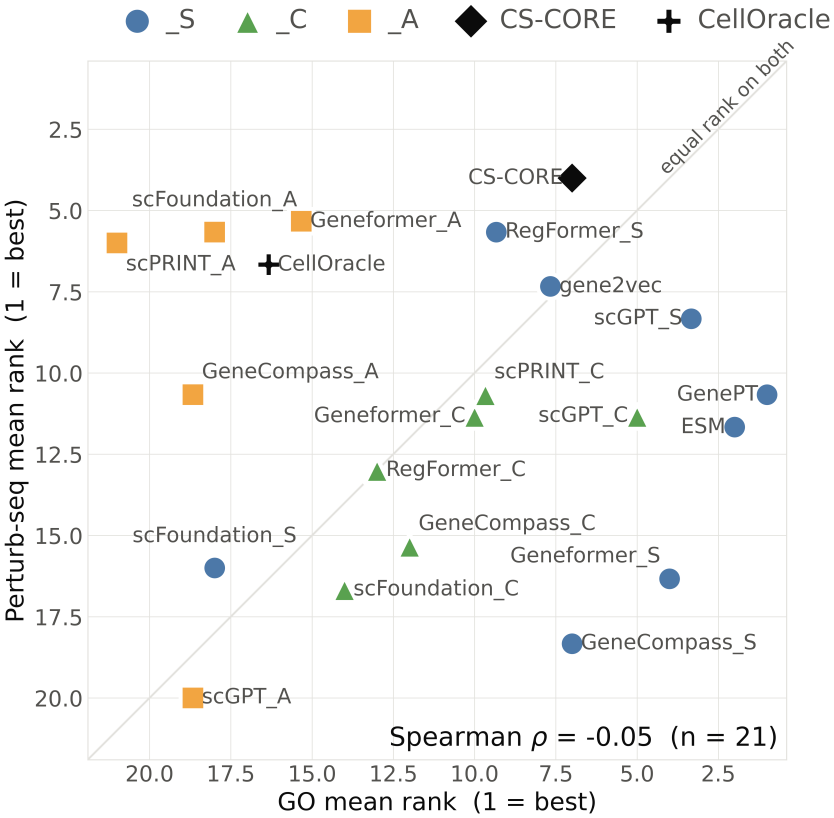
Method comparison by GO and Perturb-seq evaluation ranks, with the diagonal indicating equal performance. Mean ranks are averaged across three metrics within each evaluation setting (GO or Perturb-seq).

Methods grounded in gene co-expression show a better balance across the two benchmarks. CS-CORE is the only method ranked within the top seven on both benchmarks (GO rank 7, perturb-seq rank 1). Gene2vec, trained on bulk co-expression, is also balanced (8th and 7th) and solid on both, though best on neither. CellOracle shows a markedly different profile: it is the strongest method on Perturb-seq DE AUROC but at chance on GO, potentially reflecting its restriction to a TF-target edge space. GenePT and ESM show the other extreme: they dominate GO but are mid-ranked on Perturb-seq. Notably, GenePT embeds curated text summaries that describe gene function from the NCBI Gene database, which are also derived from literature as GO; and ESM’s protein-sequence similarity captures domain and family homology, which are often associated with shared GO annotations. By construction, neither incorporates context-specific information on gene associations.

Averaging each method’s GO and Perturb-seq mean ranks yields a single overall score. The top third (7 of 21 methods) comprises CS-CORE (5.5), GenePT (5.8), scGPT_S (5.8), ESM (6.8), RegFormer_S (7.5), gene2vec (7.5), and scGPT_C (8.2). These methods draw on sub-stantially different sources of gene relationships, including literature-derived knowledge, single-cell pretraining, observational single-cell expression, and protein-sequence information. This diversity suggests that no single representation paradigm consistently dominates GN inference. Notably, the conventional gene2vec representation ranks ahead of several scFM representations and association scores, while GRN-aware scFMs do not consistently outperform general-purpose models: RegFormer_S enters the top third, but Reg-Former_C and both scPRINT readings do not. Overall, scFM representations and association scores can achieve leading performance under specific evaluation criteria but their advantage is not universal. Their performance remains competitive with, rather than consistently superior to, conventional representation learning method gene2vec and gene relationships inferred directly with domain-specific tools.

### 3.3 Context-independent versus context-dependent representations and association scores

To assess how dependence on the target dataset affects GN inference, we grouped the evaluated representations and association scores into context-independent and context-dependent groups. The context-independent group contains eight static embeddings, in which each gene is represented by a fixed vector shared across datasets. The context-dependent group contains eleven representations or association scores derived from the target dataset, including contextual embeddings and attention scores. CS-CORE, which is the domain-specific method showing relatively strong performance across metrics, was analyzed separately as a domain-specific baseline inferred directly from the target dataset. Multiple-testing correction was performed using the Benjamini-Hochberg procedure across the six comparisons shown in Figure 5, and significance was indicated by an asterisk (two-sided Wilcoxon signed-rank tests, FDR-adjusted p-value < 0.05).

**Figure 5:**
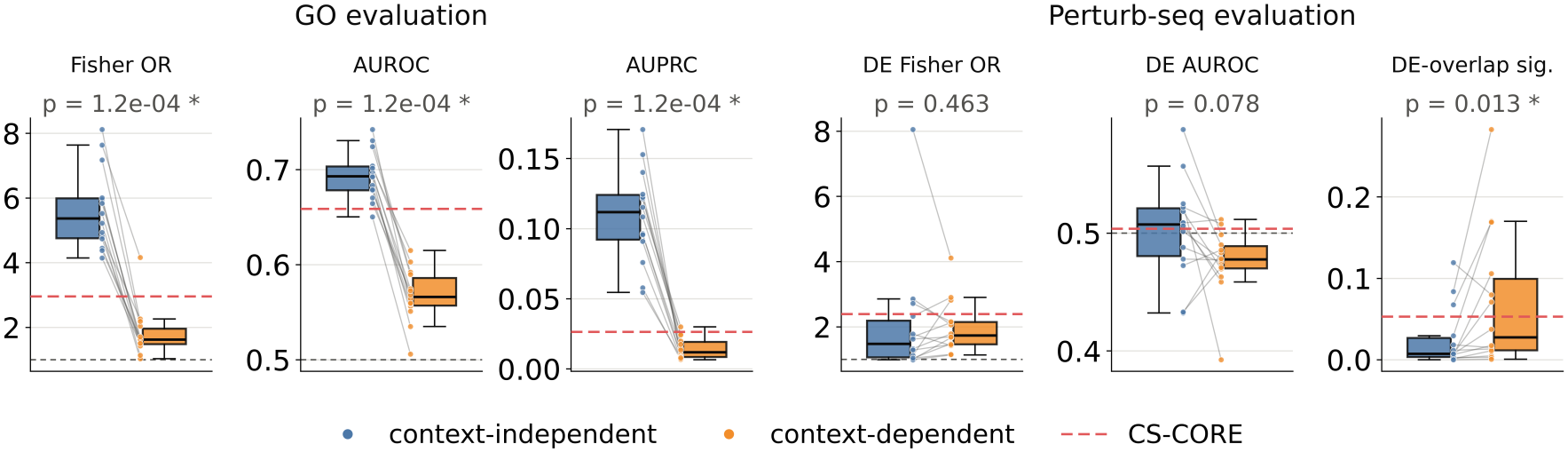
Comparison of context-independent and context-dependent representations or association scores across functional and perturbation-based metrics. For each dataset, each point shows the mean performance across gene representations or association scores within the corresponding group. Paired comparisons use two-sided Wilcoxon signed-rank tests. Means from the same dataset are connected by lines and summarized by boxplots. Gray dashed lines indicate random baselines, whereas red dashed lines denote CS-CORE.

Across all three GO-based metrics, context-independent representations consistently outperformed context-dependent representations and association scores across all 14 datasets (ΔAUROC = 0.13), indicating stronger alignment with GO-defined functional associations.

In contrast, the perturbation-based metrics revealed a divergence between edge-set enrichment and edge-ranking performance. DE Fisher OR and regulator-level enrichment were higher for context-dependent representations and association scores, with the latter showing a 3.8-fold higher median and reaching statistical significance, indicating stronger recovery of perturbation-associated DE relationships. However, DE AUROC favored context-independent representations, though it is not statistically significant.

Taken together, these results highlight that context-independent representations consistently outperform in GO evaluation while context-dependent representations and association scores show advantage on certain Perturb-seq metrics. As method performance depends on both the reference evidence (GO or Perturb-seq) and the evaluation metric, no representation consistently performs well across all evaluation settings.

### 3.4 Robustness, Sensitivity, and Computational Cost

GNEval rankings were robust to network size, thresholds for building GO-based evaluation, and the number and population of cells used for contextual extraction, with mean pairwise Spearman correlations of 0.94 across network sizes, 0.90 across 250-4,000-cell subsamples, and 0.88 across mixed versus control-only input cells (Appendix J.1, Appendix Figure 6). GO-based rankings were similarly stable across thresholds (*ρ* = 0.91), supporting the robustness of the main conclusions.

GNEval performance showed no significant systematic differences across tissues, cell systems, or perturbation-modality groups after multiple-testing correction (Appendix J.2, Appendix Table 13), suggesting that the observed patterns were not primarily driven by these dataset characteristics.

Finally, scFM embedding and attention score extraction remained computationally tractable at the evaluated scale, with contextual methods benchmarked on a single NVIDIA A30 GPU and domain-specific methods on eight CPU cores (Appendix Figure 7). Computational cost was therefore not a primary bottleneck of GN inference task.

## 4 Discussion, Conclusion and Limitations

Our benchmark results show that scFMs can be useful for gene network inference in small-sample settings. By leveraging knowledge acquired during large-scale pretraining, scFMs can recover biologically meaningful gene relationships and complement domain-specific methods. However, their utility depends on the type of representation or association score and the biological relationships of interest.

Our study highlights three main takeaways. First, gene representations should be selected in a task-aware manner rather than by a single leaderboard: context-independent representations better recover GO-defined functional relationships, whereas context-dependent representations better capture regulator-level target-set enrichment. Second, model rankings are highly sensitive to how gene-network quality is defined and evaluated. Perturbation performance remains metric-dependent, and, notably, rankings across the GO- and Perturb-seq-based benchmarks are nearly uncorrelated. Third, the distinct strengths of CS-CORE, gene2vec, and CellOracle across different metrics further motivate task-aware method selection and suggest potential benefits from integrating scFMs with domain-specific approaches.

### Broader implications

At its core, our work addresses a general problem: inferring the dependency structure among features from a small dataset, using a foundation model pretrained on large-scale data as a transferable source of prior knowledge. This problem arises in many domains. In neuroimaging, functional connectivity networks among brain regions are typically estimated from clinical cohorts of tens of subjects, while foundation models for brain activity are now pretrained on thousands of recordings (e.g., Ortega Caro et al., 2024). In multivariate time series, dependency graphs among sensors, assets, or climate variables are often inferred from short records, while time-series foundation models are pretrained on massive corpora (e.g., Ansari et al., 2024; Das et al., 2024). In each case, the same question arises: can the relationship between features encoded in a foundation model improve network inference on a small new dataset? Single-cell biology is a particularly suitable testbed for these questions, because Perturb-seq provides large-scale interventional evidence for validating inferred networks, and Gene Ontology database curate existing literature on gene-gene relationship, which are rarely available in other domains. Our study exemplifies how this broader question can be investigated by systematically evaluating the gene relationships encoded in pretrained foundation models against complementary biological evidence and comparing their performance with domain-specific methods.

### Limitations

GNEval has several limitations. The analysis is restricted to approximately 1,000 highly variable genes, which improves computational tractability but narrows the evaluated gene space and may underrepresent lowly expressed regulators. Perturbation-derived DE evidence may be limited by statistical power, and only experimentally tested regulators can be evaluated. In addition, overlap between public model pretraining corpora and benchmark datasets cannot be fully excluded, although recent Perturb-seq datasets were preferentially included to reduce this risk. The strong GO performance of GenePT may also partly reflect semantic overlap between biomedical text and ontology annotations. To reflect the practical use of scFMs in resource-limited settings, we evaluate gene representations and associations in a zero-shot setting without computationally intensive fine-tuning. While this design improves accessibility, it may not capture the full potential of fine-tuned scFM models.

### Future work

Future work will extend GNEval to larger gene sets and broader biological contexts, evaluate fine-tuned scFMs, and further integrate domain-specific network inference approaches.

### Other types of networks

This work focuses on undirected association networks, commonly known as co-expression networks, rather than directed regulatory relationship. Inferring directed regulatory relationships generally requires additional assumptions or complementary data and represents an important direction for future work.

## Data and code availability

All datasets used in this study are publicly available from their original sources. Code and scripts for data processing, representation extraction, network construction, model evaluation, and down-stream analysis are available at https://github.com/tianxinghu/GNEval-benchmark.

## Acknowledgment

This work was supported by the Rollins School of Public Health High Performance Computing Cluster at Emory University.

