## Supplementary Materials for "GNEval: A Context-Specific Benchmark for Gene Network Inference with Single-Cell Foundation Models"

### Appendix A Complete dataset description

Table 3: Additional information on the Perturb-seq datasets used for evaluation. "Cells $\times$ genes" reports the input matrix size; "Response genes" denotes the per-dataset response universe, consisting of the top 1,000 highly variable genes plus force-included targeted regulators; "Tested Regulators" is the number of regulators retained after filtering; "ctrl cells" is the number of non-targeting control cells; and "Sig. DE edges" is the number of significant regulator-gene pairs identified by SCEPTRE (BH-FDR $< 0.05$ and $|\log_2 \mathrm{FC}| > 0.25$).

| Data | Perturbation | Cell type | Cells × genes | Response genes | Tested Regulators | ctrl cells | Sig. DE edges |
|---|---|---|---|---|---|---|---|
| Adamson et al. (2016) | CRISPRi | K562 | 65,145 × 32,643 | 1083 | 87 | 7,629 | 8,386 |
| Norman et al. (2019) | CRISPRa | K562 | 69,686 × 33,694 | 1073 | 102 | 11,855 | 9,775 |
| Replogle et al. (2022) | CRISPRi | K562 | 310,385 × 8,563 | 1000 | 135 | 10,691 | 11,781 |
| Replogle et al. (2022) | CRISPRi | RPE1 | 247,914 × 8,749 | 1000 | 150 | 11,485 | 24,615 |
| Schmidt et al. (2022) | CRISPRa | primary T cells | 61,041 × 36,601 | 1020 | 72 | 4,000 | 6,220 |
| Sunshine et al. (2023) | CRISPRi | Calu-3 | 35,729 × 33,549 | 1174 | 174 | 3,541 | 1,334 |
| Otto et al. (2023) | CRISPRko | MOLM-13 | 31,069 × 36,601 | 1023 | 24 | 1,367 | 1,244 |
| Schnitzler et al. (2024) | CRISPRi | endothelial | 214,449 × 36,601 | 1000 | 72 | 5,506 | 98 |
| Tian et al. (2021) | CRISPRi | iPSC neuron | 32,300 × 33,514 | 1177 | 181 | 437 | 143 |
| Tian et al. (2021) | CRISPRa | iPSC neuron | 21,193 × 33,514 | 1081 | 95 | 434 | 999 |
| Dräger et al. (2022) | CRISPRi | iPSC microglia | 10,377 × 33,514 | 1035 | 36 | 817 | 1,375 |
| Leng et al. (2022) | CRISPRi | iPSC astrocyte | 7,933 × 33,514 | 1025 | 28 | 273 | 168 |
| Zhu et al. (2026) | CRISPRi | primary CD4 T cells | 7,282,273 × 18,130 | 1000 | 680 | 309,554 | 51,311 |
| Shi et al. (2026) | CRISPR (*in vivo*, mouse) | excitatory neuron | 2,246,807 × 16,684 | 1000 | 180 | 199,649 | 186 |

### Appendix B Data preprocessing

#### B.1 Overview

We apply two principles consistently across all datasets:

1. **Single perturbations only.** Combination and multi-gene perturbations are excluded.

2. **Low-MOI representation with non-targeting controls.** A control cell is defined as a cell whose detected guides are all non-targeting. Cells with no detected guide are excluded rather than treated as controls.

#### B.2 Common preprocessing pipeline

For each dataset, SCEPTRE DE testing, GN inference, and downstream evaluation are all restricted to the same filtered gene universe. (Peidli et al., 2024; Wei et al., 2025; Yang et al., 2025; He et al.,

2025)

##### B.2.1 Guide calling and single-perturbation selection

The datasets differ in the extent to which guide assignments are provided by the original authors. We therefore use three guide-processing strategies, all of which produce the same final representation: one retained cell associated with at most one perturbation target.

**Author-provided per-cell calls.** For datasets with author-provided perturbation assignments, we use those calls directly and retain only unambiguous single-guide or singlet cells. Cells annotated as multiplets, ambiguous assignments, or combinatorial perturbations are excluded.

**Guide-capture UMI tables without per-cell calls.** For datasets providing only guide-by-cell UMI counts, perturbation assignments are inferred using a shared procedure. A guide is considered present only if it has at least 3 UMIs, duplicate cell-guide entries are collapsed by retaining the maximum UMI count, and a cell is retained only if exactly one guide remains after filtering. Cells with no qualifying guide or multiple qualifying guides are excluded, and retained barcodes are intersected with the dataset's filtered cell set.

**High-MOI screens reduced to low-MOI subsets.** For high-MOI screens, we retain only cells compatible with a single-perturbation representation and satisfying the available singlet-quality criteria. Cells targeting multiple genes are excluded, whereas cells containing only non-targeting guides are retained as controls. When multiple guides target the same gene, they are collapsed into a single perturbation for downstream SCEPTRE analysis.

##### B.2.2 Gene and cell quality control

Quality control is applied to the raw count matrix produced by each dataset adapter. Duplicated gene symbols are removed first, because duplicated symbols cannot be mapped unambiguously to gene embeddings, GO annotations, or SCEPTRE responses. The remaining QC steps are applied in the following order:

1. **filter_genes(min_cells = 3).** Genes detected in fewer than 3 cells are removed.

2. **filter_cells(min_genes = 200).** Cells expressing fewer than 200 genes are removed.

3. **Depth covariates.** Before any gene subsetting, each cell's total UMI count and number of detected genes are recorded from the full transcriptome.

4. **Mitochondrial fraction.** The mitochondrial UMI fraction is computed over the full gene set using genes matching ^MT- (case-insensitive). Cells above the threshold are removed. The default threshold is 0.1; it is increased to 0.2 for Replogle_K562 and 0.25 for Schmidt because, in these datasets, a threshold of 0.1 falls within the main body of the observed distribution rather than its upper tail.

5. **Depth quantile clipping.** Cells outside the $[0.01, 0.99]$ quantile range of either depth covariate are removed, matching the intent of SCEPTRE's response_n_umis_range and response_n_nonzero_range defaults. These internal SCEPTRE filters are disabled because they would otherwise be applied after variable-gene selection. Both QC masks are computed on the original distribution and then combined, avoiding sequential clipping of an already-trimmed population.

##### B.2.3 Normalization and variable-gene selection

Counts from retained cells are normalized to $10^4$ UMIs per cell and $\log(1 + x)$-transformed, after which the top 1,000 highly variable genes are selected using the Seurat procedure. Perturbation target genes are force-included when feasible, so the response universe consists of the top-1,000 highly variable genes together with QC-passing targets. For screens with more than 500 targets, force-inclusion is omitted for computational tractability, and regulators are restricted to targets overlapping the top-1,000-gene universe.

##### B.2.4 Test grid supplied to sceptre

The response set is the gene universe defined in Section B.2.3. Raw counts are supplied to SCEPTRE; the normalized expression matrix is used only for variable-gene selection, whereas SCEPTRE fits its own negative-binomial model to the count data. Regulators are restricted to perturbation targets that belong to the response universe and have at least **20** retained cells after QC. Cells with zero counts across all response genes are excluded because the model includes the log response depth.

The discovery set is the full regulator $\times$ response grid. Covariates include the two log-transformed depth measures defined in Section B.2.2 and, for datasets assembled from multiple 10x libraries or samples, a batch factor derived from sample identity.

##### B.3 Dataset-specific preprocessing

All datasets follow the shared preprocessing pipeline described above. Table 4 summarizes only dataset-specific processing decisions.

Table 4: Dataset-specific preprocessing of the Perturb-seq datasets used for evaluation.

| Dataset | Accession | Dataset-specific preprocessing |
|---|---|---|
| Adamson et al. (2016) | GSE90546 | Only the largest single batch (approximately 65k cells) is used; the three available batches are highly imbalanced and are not combined. |
| Norman et al. (2019) | GSE133344 | Cells containing only non-targeting guides are treated as controls. Cells targeting exactly one gene are retained, whereas two-gene combinatorial perturbations are excluded. |
| Replogle et al. (2022) | Figshare 20029387 | The authors' per-cell guide and target assignments are used directly, with non-targeting guides mapped to NT. Each retained cell carries a single guide. |
| Replogle et al. (2022) | Figshare 20029387 | The authors' per-cell guide and target assignments are used directly, with non-targeting guides mapped to NT. Each retained cell carries a single guide. |
| Schmidt et al. (2022) | GSE190604 | Only single-guide cells are retained, and non-targeting guides are mapped to NT. |
| Sunshine et al. (2023) | GSE208240 | Cells without a guide assignment are excluded. Non-targeting-only cells are controls, single-target cells are retained, and multi-target cells are excluded to obtain a low-MOI-compatible subset. |
| Otto et al. (2023) | GSE200201 | Only the single-guide RNA-seq samples are used, and cells are retained only when assigned exactly one guide. |
| Schnitzler et al. (2024) | GSE210681 | Only high-confidence single-guide cells are retained. Non-targeting and safe-targeting controls are mapped to NT. |
| Tian et al. (2021) | GSE152988 | The four CRISPRi libraries are analyzed separately from CRISPRa. Author-provided single-guide assignments are used directly, non-targeting guides are mapped to NT, and library identity is included as the batch covariate. |
| Tian et al. (2021) | GSE152988 | The two CRISPRa libraries are analyzed separately from CRISPRi. Author-provided single-guide assignments are used directly, non-targeting guides are mapped to NT, and library identity is included as the batch covariate. |
| Dräger et al. (2022) | GSE178317 | Author-provided per-cell guide assignments are used, retaining only cells classified as singlets. Non-targeting controls are mapped to NT. |
| Leng et al. (2022) | GSE182308 | Only the two vehicle-condition wells are used. Guide assignments are inferred from guide-capture UMI counts using the shared UMI-based procedure, and non-targeting guides are mapped to NT; well identity is used as the batch covariate. |
| Zhu et al. (2026) | — | Only resting-condition, single-guide cells are retained across the four donors, with donor identity used as the batch covariate. Because of the dataset size, cell QC and highly variable gene selection are performed using a streaming implementation equivalent to the shared pipeline. |
| Shi et al. (2026) | — | Only excitatory neurons with exactly one guide and a singlet call are retained. Non-targeting and safe-targeting guides are mapped to NT. Mouse genes and targets are mapped to human orthologs, and the large expression matrix is processed using the streaming implementation of the shared pipeline. |

### Appendix C Differential-expression reference

We constructed the perturbational reference for each dataset using SCEPTRE (Barry et al., 2024) (v0.10.3). All differential-expression tests were performed on raw UMI counts; normalized and log-transformed expression values were used only for highly variable gene selection (Section B.2.3).

#### C.1 SCEPTRE analysis

We analyzed all datasets under the low-MOI setting. Perturbation assignments were taken from the single-guide calls defined in Section B.2.1; guides targeting the same gene were combined using the SCEPTRE `union` strategy.

Tests were adjusted for log-transformed sequencing depth and detected-gene count, both computed from the full transcriptome before gene selection. A sample or library batch covariate was additionally included where applicable. Because cell-level QC had already been performed using the full transcriptome (Section B.2.2), SCEPTRE's response-matrix-based cellwise QC thresholds were disabled. We retained its pairwise sparsity filter, requiring at least seven cells with nonzero expression of the response gene in both the treatment and non-targeting control groups.

All regulator-response pairs in the analysis universe were tested using two-sided low-MOI SCEPTRE tests. We additionally ran SCEPTRE's negative-control calibration check for each dataset.

#### C.2 Definition of perturbational edges

We did not use SCEPTRE's default significance calls. Instead, $p$-values were adjusted by the Benjamini-Hochberg procedure separately within each regulator across its tested response genes. A regulator–response pair $(r, g)$ was considered a DE edge when

$$q_{rg} < 0.05 \qquad \text{and} \qquad |\log_2 \mathrm{FC}_{rg}| > 0.25.$$

Self-pairs $(r = g)$ were excluded.

Because the inferred networks evaluated in this study are undirected and unsigned, DE edges were converted to unordered gene pairs irrespective of effect direction or sign. Evaluation was restricted to the testable subgraph: pairs for which at least one endpoint was a regulator subjected to perturbational testing. Pairs between two non-regulators were therefore treated as untested rather than as negative examples.

### Appendix D Representation extraction

For each method, we use the released checkpoint and follow the original representation-extraction procedure whenever available. Table 5 summarizes the checkpoint, readout, aggregation, dimensionality, and coverage of every representation.

**Shared conventions.** All representations are aligned to the dataset-specific gene universe defined in Appendix B. Genes without a valid representation are flagged and excluded from downstream network construction rather than treated as zero vectors. Contextual and attention representations use the same cells per dataset: all post-QC non-targeting control cells (capped at 12,000 for the two largest screens), the cells CS-CORE is fit on. Sample-size experiments use nested subsets of 250-4,000 cells. Static representations require no cell input.

For attention-based networks, gene-gene scores are symmetrized by averaging the two directions. When a model tokenizes only expressed genes, each pair is averaged over cells in which both genes are present, and pairs observed in fewer than 10 cells are discarded. Embeddings are otherwise used as extracted, with no additional centering, whitening, or normalization before cosine similarity.

#### D.1 Static gene representations

**GenePT.** We use the released GenePT v2 gene embeddings (Chen and Zou, 2025), generated by applying `text-embedding-ada-002` to NCBI RefSeq gene summaries. The released 1,536-dimensional vectors are used directly, with no additional preprocessing.

**gene2vec.** We use the released `gene2vec_dim_200_iter_9.txt` embeddings (Du et al., 2019), trained by skip-gram on gene co-expression relationships derived from bulk GEO data. The 200-dimensional vectors are used as published.

**ESM-2.** We use the gene-level ESM-2 15B representations distributed with UCE (Lin et al., 2023; Rosen et al., 2026). Protein representations are obtained from the final ESM-2 layer, mean-pooled

Table 5: Representations used in the benchmark. **S**: static embedding; **C**: contextual embedding; **A**: attention. Coverage is the median fraction of the dataset-specific gene panel represented across the evaluated datasets.

| **Method** | **Checkpoint** | **Layer / tensor** | **Aggregation** | **Dim** | **Coverage** |
|---|---|---|---|---|---|
| GenePT (S) | GenePT v2, `ada_text` | text-encoder output | — | 1536 | 84.0% |
| gene2vec (S) | `dim_200_iter_9` | word2vec vector | — | 200 | 80.5% |
| ESM-2 (S) | ESM-2 15B (via UCE) | final repr. layer | mean residues, then isoforms | 5120 | 77.9% |
| scGPT (S) | `scGPT_human` | token emb. + `enc_norm` | — | 512 | 87.9% |
| scGPT (C) | `scGPT_human` | layer 12 hidden state | mean over cells | 512 | 87.9% |
| scGPT (A) | `scGPT_human` | layer 12 $q, k$ | mean over heads/cells | — | 87.9% |
| Geneformer (S) | V2-104M | input token emb. table | — | 768 | 74.6% |
| Geneformer (C) | V2-104M | layer 12 hidden state | mean over detecting cells | 768 | 64.3% |
| Geneformer (A) | V2-104M | layer 12 attention | mean over heads/co-detecting cells | — | 64.3% |
| scFoundation (S) | `models.ckpt` | `pos_emb` table | — | 768 | 74.7% |
| scFoundation (C) | `models.ckpt` | decoder output | mean over cells | 512 | 74.7% |
| scFoundation (A) | `models.ckpt` | encoder layer 12 attention | mean over heads/co-expressing cells | — | 64.3% |
| GeneCompass (S) | GeneCompass_Base | word-emb. table | — | 768 | 77.1% |
| GeneCompass (C) | GeneCompass_Base | layer 12 hidden state | mean over expressing cells | 768 | 66.3% |
| GeneCompass (A) | GeneCompass_Base | layer 12 attention | mean over heads/co-expressing cells | — | 66.3% |
| RegFormer (S) | RegFormer-10k | encoder emb. table | — | 512 | 87.9% |
| RegFormer (C) | RegFormer-10k | final Mamba block | mean over expressing cells | 512 | 63.1% |
| scPRINT (C) | `medium-v1.5` | final block gene emb. | mean over expressing cells | 256 | 66.2% |
| scPRINT (A) | `medium-v1.5` | layer 8 shared-QK | official GNInfer aggregation | — | 66.2% |
| CS-CORE | — | test statistic | all control cells ($\leq$12,000) | — | 100% |

over residues and then averaged across protein isoforms to obtain one vector per gene.

**scGPT.** For the static scGPT representation (Cui et al., 2024), we read the gene encoder from the released `scGPT_human` checkpoint. The representation is the token-embedding table followed by the encoder's own layer normalization.

**Geneformer.** Static Geneformer representations (Theodoris et al., 2023) are read directly from the V2-104M input embedding table. Special tokens are removed and Ensembl identifiers are mapped to gene symbols using the released dictionary.

**scFoundation.** For scFoundation (Hao et al., 2024), the static representation is the released `pos_emb` gene-identity table aligned to the model's fixed 19,264-gene vocabulary.

**GeneCompass.** Static GeneCompass representations (Yang et al., 2024) are taken from the GeneCompass_Base word-embedding table, restricted to human gene tokens and mapped to gene symbols using the repository-provided dictionary.

**RegFormer.** Static RegFormer representations (Hu et al., 2026) are taken directly from the RegFormer-10k gene-token embedding table. Its vocabulary matches that of the scGPT whole-human checkpoint.

#### D.2 Contextual representations and attention

**scGPT.** Raw counts from the shared control-cell set are processed with scGPT's `Preprocessor`: library-size normalization to $10^4$, $\log(1 + x)$ transformation, and 51-bin discretization. The shared gene panel is retained rather than re-selecting HVGs. Contextual gene embeddings are the final-layer hidden states averaged across cells. Attention is reconstructed from the final layer's queries and keys using the model's released gene network procedure, then averaged across heads and cells.

**Geneformer.** Cells are tokenized with the official V2 `TranscriptomeTokenizer`, which ranks detected genes after normalization by pretraining median expression. Contextual representations are final-layer hidden states averaged over cells in which each gene is detected. Attention uses the final-layer post-softmax attention and is averaged over heads and co-detecting cells.

**scFoundation.** Inputs follow the released inference workflow: $\log(1 + x/\text{total} \times 10^4)$ on the model's fixed gene axis together with the two resolution tokens used during pretraining. Contextual representations are taken from the decoder output and averaged across cells. Attention is taken from the final encoder layer and averaged across heads and cells in which both genes are expressed.

**GeneCompass.** GeneCompass inputs consist of nonzero genes ordered by normalized expression using the model's released preprocessing. Contextual representations are final-layer hidden states averaged across cells expressing each gene. Attention is the model's final-layer softmax attention, averaged over heads and co-expressing cells. We follow the repository's released embedding-generation procedure for loading its prior-knowledge embeddings.

**RegFormer.** We use the repository's `infer_dataset` pipeline with its released preprocessing and graph-based token ordering. Contextual gene embeddings are taken from the final Mamba block and averaged over cells in which each gene is expressed. RegFormer provides no attention representation because its encoder uses Mamba blocks rather than self-attention.

**scPRINT.** Raw counts are aligned to the checkpoint's human gene axis and passed through the released scPRINT inference pipeline (Kalfon et al., 2025). Contextual embeddings are the final block's gene representations averaged over expressing cells. Attention-based networks use the official `GNInfer` shared-query/key procedure, which averages queries and keys across cells before computing the gene–gene score.

##### D.3 CS-CORE baseline

CS-CORE (Su et al., 2023) is fit directly to raw UMI counts from all post-QC non-targeting control cells (capped at 12,000 for the two largest screens, as for the contextual representations) on the shared gene panel. The signed CS-CORE test statistic is used directly as the gene-gene edge score, without an intermediate embedding or similarity calculation.

##### D.4 CellOracle baseline

CellOracle (Kamimoto et al., 2023) infers directed TF-target edges by combining a promoter motif-based prior with bagging ridge regression on imputed single-cell expression, and retains significant edges ranked by the absolute regression coefficient. We reproduce this procedure on all control cells of each dataset, using the same significance filtering and edge ranking, then symmetrize directed edges by keeping the stronger direction and evaluate the resulting network at the available sizes.

#### Appendix E Network construction

Each representation in Appendix D is converted into an undirected gene network by defining a gene-gene score, restricting the gene space to genes represented by the method, and selecting the strongest edges.

##### E.1 Gene-gene scores

For static and contextual gene embeddings, edge scores are cosine similarities:

$$S_{ij} = \frac{\mathbf{x}_i^\top \mathbf{x}_j}{\|\mathbf{x}_i\| \, \|\mathbf{x}_j\|}.$$

Representations are used as extracted. Attention matrices and CS-CORE already provide gene-gene scores and are therefore used directly without an additional similarity calculation.

Alternative similarity constructions, including principal-component residualization, rank-normalized cosine similarity, and graphical lasso, are evaluated separately as robustness analyses.

#### E.2 Gene space and network conventions

For each method and dataset, the network is constructed on the intersection between the dataset's response-gene universe and the genes for which that method provides a valid representation. Genes outside this intersection are not imputed or treated as zero vectors.

Self-pairs are excluded. All evaluated networks are undirected; asymmetric attention matrices are symmetrized as

$$\tilde{A} = \frac{1}{2}(A + A^{\top}).$$

For signed scores, edge selection uses absolute magnitude, $|S_{ij}|$.

#### E.3 Edge selection and network size

For a target network size $K$, we select the $K$ gene pairs with the largest absolute scores from the strict upper triangle:

$$E_K = \mathrm{TopK}_{i<j}\{|S_{ij}|\}.$$

This rank-based construction allows representations with different numerical score scales, including cosine similarity, attention, and CS-CORE statistics, to be compared at the same edge count.

The main benchmark evaluates

$$K \in \{5{,}000,\ 10{,}000,\ 20{,}000\},$$

with reported results summarized across these network sizes.

Exact-zero or otherwise unavailable gene pairs are not promoted to edges. All methods except CellOracle have sufficient valid candidate pairs to reach the three network sizes used in the main benchmark.

Unlike the other methods, CellOracle produces sparse networks restricted to TF-target pairs supported by its base GRN, which is inferred from external data. Following the recommended filtering procedure ($p < 0.001$), its networks contain 1,430-12,767 edges across datasets. CellOracle is therefore evaluated at each required network size (5,000, 10,000 and 20,000 edges) that its network reaches, together with its full network, and is excluded from sizes it does not reach; on the four datasets whose full network has fewer than 5,000 edges, it is evaluated on the full network alone.

### Appendix F GO-based evaluation

We evaluate whether predicted gene pairs share functional annotation using the gene-to-GO mapping distributed with GEARS (Roohani et al., 2023). The same fixed annotation snapshot is used for all datasets and methods.

#### F.1 GO reference

For each dataset, let $\mathcal{G}$ denote its response-gene universe and $\mathrm{GO}(g)$ the set of GO terms annotated to gene $g$. For every annotated gene pair, we compute the Jaccard similarity

$$J(g_1, g_2) = \frac{|\mathrm{GO}(g_1) \cap \mathrm{GO}(g_2)|}{|\mathrm{GO}(g_1) \cup \mathrm{GO}(g_2)|}.$$

A pair is considered a GO-positive if $J > \tau$. We use

$$\tau \in \{0.1,\ 0.3\},$$

with $\tau = 0.3$ as the primary reference in the main analysis and $\tau = 0.1$ as a less stringent robustness analysis. Genes absent from the annotation source are excluded from the GO reference.

#### F.2 Evaluation metrics

For each predicted network, AUROC and AUPRC are computed over the selected edges, using $|S_{ij}|$ as the ranking score and GO membership as the binary label. AUROC has chance level 0.5; for AUPRC we additionally record the positive prevalence among the selected edges. A network with no GO-positive pair among its selected edges is assigned an AUROC of 0.5 and an AUPRC of 0, as for the perturbational metrics.

Set-level enrichment is measured with a one-sided Fisher exact test comparing selected versus unselected pairs and GO-positive versus GO-negative pairs. The reported effect size is the Fisher odds ratio.

Because methods differ in gene coverage, the Fisher background is defined over the dataset's full response-gene universe rather than each method's represented gene subset. This keeps the reference prevalence fixed across methods within a dataset. AUROC and AUPRC are unaffected by this background choice because they are computed over selected edges.

### Appendix G Perturbational evaluation

We evaluate whether predicted gene pairs are supported by perturbation-induced differential expression. Unlike the GO evaluation, this reference is based on experimental intervention: one gene is perturbed and the transcriptional response of other genes is measured.

#### G.1 Differential-expression reference

For each dataset, tested regulators are perturbation targets that belong to the response-gene universe and have at least 20 retained perturbed cells. Each regulator is tested against all response genes using SCEPTRE (Barry et al., 2024) on raw UMI counts.

Tests are two-sided and use non-targeting cells as controls where available. Covariates include log sequencing depth, log detected-gene count, and batch where applicable; depth covariates are computed from the full transcriptome before gene selection. A regulator-response pair is tested only when the response gene is detected in at least seven treatment and seven control cells.

A pair is called differentially expressed when

$$q < 0.05 \qquad \text{and} \qquad |\log_2 \mathrm{FC}| > 0.25,$$

where Benjamini-Hochberg correction is applied separately within each regulator. Self-pairs are excluded.

Because the evaluated networks are undirected, a significant (regulator, response) result is converted to the unordered pair $\{r, g\}$.

#### G.2 Testable pair space

Perturbational evaluation is restricted to the *testable subgraph*: gene pairs for which at least one endpoint is a tested regulator. Pairs between two non-regulators cannot be supported by the perturbation experiment and are therefore treated as untested rather than as negatives.

For a response universe of size $p$ and regulator set $R$, the testable pair space contains

$$N_t = \binom{p}{2} - \binom{p - |R|}{2}$$

unordered pairs.

#### G.3 Evaluation metrics

We evaluate perturbational agreement at three complementary levels.

**Set-level enrichment.** Predicted testable edges are compared with DE-supported pairs using a one-sided Fisher exact test over the testable subgraph. We report the odds ratio, with an odds ratio of 1 corresponding to random enrichment.

**Edge-level ranking.** AUROC is computed over predicted testable edges, using the network edge score as the ranking variable and DE support as the binary label. Chance performance is 0.5.

**Regulator-level enrichment.** For each tested regulator $r$, we compare its predicted neighborhood $\mathcal{N}(r)$ with the set of genes $D(r)$ that respond significantly to perturbation of $r$. A one-sided Fisher test is performed for each regulator, followed by BH correction across regulators. We report

$$\text{enrichment rate} = \frac{\#\{r : \text{FDR}(r) < 0.05\}}{\#\{r \text{ tested}\}},$$

the fraction of regulators whose predicted neighborhoods are significantly enriched for their measured transcriptional responses.

##### G.4 Filtering and aggregation

A network with no DE-supported pair among its testable edges is assigned an odds ratio of 0 and an AUROC of 0.5 rather than being dropped, and regulator-level evaluation requires at least one tested regulator, so every method is scored on all 14 datasets.

For each metric, values are first averaged across the three network sizes within each method-dataset pair. Method groups are then compared across datasets using two-sided paired Wilcoxon signed-rank tests. Multiple comparisons are corrected using Benjamini-Hochberg where applicable.

##### G.5 Interpretation

These metrics provide complementary views of perturbational agreement: the Fisher odds ratio measures whether the predicted edge set is enriched for perturbation-supported pairs, AUROC measures whether the network ranks such pairs highly, and the regulator-level enrichment rate measures whether individual predicted neighborhoods recover experimentally responsive genes.

#### Appendix H Statistical analysis

The dataset is the unit of statistical inference throughout. For each method-dataset pair, metrics are first averaged across the three network sizes. For representation-class comparisons, method-level values are then averaged within each comparison group, yielding one paired value per dataset. Methods and network sizes are therefore not treated as independent replicates.

Paired comparisons, including context-independent versus context-dependent representations, use two-sided Wilcoxon signed-rank tests. We report the median paired difference and matched-pairs rank-biserial correlation,

$$r = \frac{W^+ - W^-}{W^+ + W^-}.$$

Comparisons between disjoint dataset groups, such as tissue, cell system, or perturbation type, use two-sided Mann-Whitney $U$ tests with Cliff's $\delta$ as the effect size. Associations between method rankings are summarized using Spearman correlation and are treated as exploratory.

Benjamini-Hochberg correction is applied separately within each predefined family of related hypothesis tests, with $q < 0.05$ considered significant. All tests are two-sided.

Because the number of datasets is small, non-significant results are interpreted as insufficient evidence for a detectable effect rather than as evidence of equivalence.

### Appendix I Per-dataset benchmark results

Tables 6–11 report the per-dataset results underlying Figures 2 and 3. Each value is averaged over networks with $K \in \{5{,}000, 10{,}000, 20{,}000\}$ edges. The final column reports the median across datasets. Rows are ordered by mean rank separately within the GO and perturbational blocks.

GO results use the Jaccard $> 0.3$ reference. Dataset abbreviations are: Adam (Adamson), Norm (Norman), K562/RPE1 (Replogle), Schm (Schmidt), SARS (Sunshine SARS-CoV-2), MOLM (Otto MOLM-13), Endo (Schnitzler endothelial), Ti-i/Ti-a (Tian2020 CRISPRi/CRISPRa), Drag (Drager2021 microglia), Leng (Leng2021 astrocyte), GWCD (GWCD4i), and WBrn (WholeBrain excitatory).

**Zero-filled cells.** Under the zero-fill rule, 11 of the 294 cells in each pair-level DE table contain no DE-supported pair at any network size and therefore read exactly 0 (odds ratio) or 0.5 (AUROC): 7 for scFoundation_C, 3 for GeneCompass_S and 1 for scGPT_S, concentrated in the low-DE-density screens. GO cells are filled the same way.

Table 6: Per-dataset GO Fisher odds ratio (Jaccard > 0.3).

| Representation | Adam | Norm | K562 | RPE1 | Schm | SARS | MOLM | Endo | Ti-i | Ti-a | Drag | Leng | GWCD | WBrn | Median |
|---|---|---|---|---|---|---|---|---|---|---|---|---|---|---|---|
| GenePT | 10.68 | 14.42 | 15.69 | 10.97 | 13.94 | 24.56 | 19.45 | 11.67 | 16.40 | 14.79 | 13.87 | 16.56 | 15.03 | 19.43 | **14.91** |
| ESM | 5.82 | 6.85 | 4.48 | 4.70 | 6.20 | 8.23 | 12.94 | 6.56 | 11.55 | 8.21 | 8.77 | 6.69 | 8.23 | 8.64 | **7.53** |
| scGPT_S | 4.20 | 3.48 | 4.02 | 5.24 | 5.16 | 3.80 | 6.41 | 4.79 | 6.83 | 3.37 | 6.64 | 6.42 | 5.47 | 2.57 | **4.97** |
| Geneformer_S | 4.75 | 4.51 | 3.22 | 3.98 | 6.16 | 6.40 | 6.10 | 4.27 | 13.70 | 5.48 | 6.30 | 5.47 | 7.73 | 4.01 | **5.47** |
| scGPT_C | 3.21 | 2.13 | 3.63 | 3.74 | 2.81 | 2.99 | 13.78 | 5.23 | 1.91 | 1.19 | 4.38 | 4.91 | 3.71 | 0.89 | **3.42** |
| GeneCompass_S | 4.49 | 4.30 | 2.31 | 3.00 | 5.54 | 5.24 | 5.34 | 2.47 | 10.68 | 3.73 | 5.10 | 4.17 | 6.01 | 3.83 | **4.39** |
| CS-CORE | 2.44 | 2.35 | 4.15 | 3.01 | 4.69 | 6.68 | 2.65 | 5.13 | 1.14 | 0.97 | 3.78 | 2.90 | 3.25 | 0.80 | **2.96** |
| gene2vec | 2.67 | 2.13 | 2.63 | 2.73 | 3.70 | 5.44 | 5.50 | 5.17 | 2.84 | 1.83 | 4.35 | 4.28 | 2.93 | 1.77 | **2.89** |
| RegFormer_S | 2.04 | 1.31 | 1.96 | 1.71 | 2.96 | 2.80 | 4.27 | 3.00 | 1.65 | 0.96 | 2.61 | 2.35 | 1.49 | 0.91 | **2.00** |
| scPRINT_C | 2.36 | 2.69 | 3.02 | 3.19 | 2.73 | 6.51 | 5.80 | 4.26 | 1.68 | 1.19 | 3.03 | 3.47 | 2.02 | 1.14 | **2.87** |
| Geneformer_C | 2.50 | 2.59 | 2.91 | 2.78 | 3.18 | 3.08 | 7.03 | 3.28 | 1.53 | 1.48 | 2.54 | 2.11 | 2.77 | 1.64 | **2.68** |
| GeneCompass_C | 1.21 | 1.34 | 1.89 | 1.97 | 1.45 | 2.18 | 3.22 | 1.91 | 2.44 | 0.86 | 1.86 | 1.55 | 1.95 | 1.62 | **1.88** |
| RegFormer_C | 2.05 | 0.98 | 1.11 | 0.78 | 1.55 | 0.92 | 6.47 | 3.15 | 1.59 | 0.87 | 1.07 | 2.36 | 1.16 | 2.03 | **1.36** |
| scFoundation_C | 1.31 | 1.04 | 0.61 | 1.08 | 0.77 | 1.79 | 1.33 | 0.54 | 10.08 | 3.36 | 0.86 | 1.19 | 3.35 | 2.30 | **1.25** |
| Geneformer_A | 1.83 | 1.63 | 0.90 | 0.76 | 1.02 | 1.96 | 0.64 | 2.14 | 0.84 | 0.53 | 0.94 | 0.91 | 0.56 | 0.33 | **0.90** |
| CellOracle | 1.27 | 1.87 | 0.95 | 1.14 | 1.01 | 1.55 | 0.86 | 0.00 | 1.02 | 2.97 | 1.14 | 0.58 | 0.89 | 1.07 | **1.05** |
| scFoundation_S | 0.93 | 0.88 | 0.72 | 0.85 | 0.51 | 0.93 | 1.06 | 0.61 | 1.30 | 1.10 | 0.39 | 0.73 | 0.65 | 0.59 | **0.79** |
| scFoundation_A | 1.25 | 0.72 | 1.01 | 0.47 | 0.37 | 0.85 | 2.07 | 1.64 | 0.79 | 0.26 | 0.49 | 0.76 | 0.40 | 1.05 | **0.77** |
| GeneCompass_A | 1.45 | 1.91 | 0.84 | 0.66 | 0.79 | 1.47 | 3.84 | 1.00 | 1.60 | 0.42 | 0.36 | 0.42 | 0.64 | 0.45 | **0.81** |
| scGPT_A | 0.61 | 0.64 | 1.23 | 0.60 | 0.77 | 0.40 | 0.69 | 0.73 | 0.73 | 0.97 | 0.63 | 0.82 | 1.05 | 0.79 | **0.73** |
| scPRINT_A | 0.43 | 0.61 | 0.72 | 0.36 | 0.27 | 0.28 | 0.92 | 0.97 | 0.84 | 0.19 | 0.44 | 0.26 | 0.29 | 0.24 | **0.39** |

Table 7: Per-dataset GO AUROC (Jaccard > 0.3).

| Representation | Adam | Norm | K562 | RPE1 | Schm | SARS | MOLM | Endo | Ti-i | Ti-a | Drag | Leng | GWCD | WBrn | Median |
|---|---|---|---|---|---|---|---|---|---|---|---|---|---|---|---|
| GenePT | 0.881 | 0.838 | 0.875 | 0.856 | 0.824 | 0.911 | 0.822 | 0.830 | 0.899 | 0.838 | 0.872 | 0.870 | 0.840 | 0.842 | **0.849** |
| ESM | 0.819 | 0.769 | 0.749 | 0.803 | 0.739 | 0.835 | 0.761 | 0.816 | 0.788 | 0.770 | 0.855 | 0.796 | 0.790 | 0.810 | **0.793** |
| scGPT_S | 0.771 | 0.749 | 0.775 | 0.708 | 0.790 | 0.807 | 0.810 | 0.842 | 0.650 | 0.599 | 0.783 | 0.740 | 0.667 | 0.642 | **0.760** |
| Geneformer_S | 0.765 | 0.687 | 0.659 | 0.634 | 0.753 | 0.805 | 0.765 | 0.633 | 0.840 | 0.659 | 0.713 | 0.670 | 0.674 | 0.711 | **0.699** |
| scGPT_C | 0.713 | 0.724 | 0.683 | 0.674 | 0.715 | 0.669 | 0.838 | 0.768 | 0.501 | 0.631 | 0.756 | 0.715 | 0.707 | 0.664 | **0.710** |
| GeneCompass_S | 0.739 | 0.638 | 0.647 | 0.591 | 0.667 | 0.737 | 0.687 | 0.532 | 0.750 | 0.620 | 0.699 | 0.642 | 0.646 | 0.667 | **0.657** |
| CS-CORE | 0.683 | 0.659 | 0.659 | 0.621 | 0.817 | 0.606 | 0.759 | 0.791 | 0.523 | 0.580 | 0.794 | 0.558 | 0.670 | 0.601 | **0.659** |
| gene2vec | 0.711 | 0.702 | 0.626 | 0.682 | 0.680 | 0.624 | 0.678 | 0.750 | 0.525 | 0.632 | 0.704 | 0.667 | 0.674 | 0.591 | **0.676** |
| RegFormer_S | 0.586 | 0.599 | 0.725 | 0.658 | 0.734 | 0.712 | 0.717 | 0.743 | 0.626 | 0.552 | 0.601 | 0.595 | 0.592 | 0.507 | **0.613** |
| scPRINT_C | 0.682 | 0.599 | 0.615 | 0.659 | 0.748 | 0.630 | 0.677 | 0.641 | 0.510 | 0.546 | 0.718 | 0.643 | 0.628 | 0.584 | **0.636** |
| Geneformer_C | 0.610 | 0.575 | 0.693 | 0.629 | 0.650 | 0.641 | 0.628 | 0.578 | 0.663 | 0.581 | 0.621 | 0.563 | 0.621 | 0.577 | **0.621** |
| GeneCompass_C | 0.664 | 0.478 | 0.559 | 0.582 | 0.671 | 0.603 | 0.572 | 0.608 | 0.602 | 0.482 | 0.656 | 0.621 | 0.646 | 0.588 | **0.603** |
| RegFormer_C | 0.693 | 0.594 | 0.660 | 0.565 | 0.627 | 0.560 | 0.692 | 0.663 | 0.519 | 0.495 | 0.597 | 0.579 | 0.566 | 0.555 | **0.587** |
| scFoundation_C | 0.602 | 0.453 | 0.641 | 0.507 | 0.487 | 0.685 | 0.541 | 0.204 | 0.644 | 0.589 | 0.536 | 0.528 | 0.591 | 0.659 | **0.565** |
| Geneformer_A | 0.677 | 0.642 | 0.574 | 0.611 | 0.437 | 0.472 | 0.455 | 0.563 | 0.469 | 0.634 | 0.669 | 0.547 | 0.465 | 0.525 | **0.555** |
| CellOracle | 0.392 | 0.537 | 0.433 | 0.403 | 0.514 | 0.603 | 0.537 | 0.500 | 0.523 | 0.606 | 0.478 | 0.354 | 0.475 | 0.466 | **0.489** |
| scFoundation_S | 0.572 | 0.485 | 0.573 | 0.491 | 0.361 | 0.506 | 0.553 | 0.427 | 0.459 | 0.533 | 0.384 | 0.450 | 0.479 | 0.545 | **0.488** |
| scFoundation_A | 0.486 | 0.472 | 0.674 | 0.381 | 0.575 | 0.414 | 0.415 | 0.713 | 0.465 | 0.262 | 0.655 | 0.551 | 0.498 | 0.622 | **0.492** |
| GeneCompass_A | 0.467 | 0.585 | 0.331 | 0.452 | 0.449 | 0.446 | 0.506 | 0.491 | 0.571 | 0.513 | 0.344 | 0.539 | 0.384 | 0.575 | **0.479** |
| scGPT_A | 0.542 | 0.404 | 0.563 | 0.608 | 0.621 | 0.646 | 0.480 | 0.534 | 0.445 | 0.456 | 0.524 | 0.464 | 0.522 | 0.524 | **0.524** |
| scPRINT_A | 0.376 | 0.534 | 0.638 | 0.578 | 0.338 | 0.378 | 0.682 | 0.442 | 0.494 | 0.378 | 0.689 | 0.405 | 0.494 | 0.425 | **0.468** |

Table 8: Per-dataset GO AUPRC (Jaccard > 0.3).

| Representation | Adam | Norm | K562 | RPE1 | Schm | SARS | MOLM | Endo | Ti-i | Ti-a | Drag | Leng | GWCD | WBrn | Median |
|---|---|---|---|---|---|---|---|---|---|---|---|---|---|---|---|
| GenePT | 0.194 | 0.226 | 0.399 | 0.301 | 0.063 | 0.361 | 0.198 | 0.083 | 0.542 | 0.220 | 0.346 | 0.376 | 0.386 | 0.415 | **0.324** |
| ESM | 0.243 | 0.163 | 0.153 | 0.226 | 0.118 | 0.399 | 0.209 | 0.228 | 0.326 | 0.200 | 0.414 | 0.369 | 0.291 | 0.319 | **0.236** |
| scGPT_S | 0.163 | 0.141 | 0.104 | 0.117 | 0.119 | 0.151 | 0.261 | 0.059 | 0.019 | 0.026 | 0.185 | 0.102 | 0.090 | 0.073 | **0.110** |
| Geneformer_S | 0.117 | 0.074 | 0.049 | 0.031 | 0.049 | 0.251 | 0.131 | 0.020 | 0.367 | 0.089 | 0.092 | 0.053 | 0.073 | 0.080 | **0.077** |
| scGPT_C | 0.023 | 0.054 | 0.099 | 0.049 | 0.031 | 0.037 | 0.185 | 0.031 | 0.010 | 0.020 | 0.067 | 0.107 | 0.108 | 0.059 | **0.051** |
| GeneCompass_S | 0.023 | 0.017 | 0.024 | 0.014 | 0.033 | 0.027 | 0.024 | 0.003 | 0.047 | 0.015 | 0.032 | 0.020 | 0.061 | 0.077 | **0.024** |
| CS-CORE | 0.014 | 0.033 | 0.055 | 0.027 | 0.034 | 0.014 | 0.026 | 0.043 | 0.004 | 0.012 | 0.068 | 0.013 | 0.034 | 0.007 | **0.026** |
| gene2vec | 0.023 | 0.060 | 0.017 | 0.030 | 0.010 | 0.017 | 0.034 | 0.019 | 0.004 | 0.020 | 0.031 | 0.040 | 0.032 | 0.016 | **0.022** |
| RegFormer_S | 0.006 | 0.046 | 0.118 | 0.045 | 0.044 | 0.016 | 0.062 | 0.049 | 0.057 | 0.033 | 0.020 | 0.017 | 0.036 | 0.005 | **0.040** |
| scPRINT_C | 0.012 | 0.013 | 0.018 | 0.020 | 0.016 | 0.022 | 0.025 | 0.010 | 0.002 | 0.004 | 0.019 | 0.017 | 0.016 | 0.081 | **0.017** |
| Geneformer_C | 0.026 | 0.010 | 0.070 | 0.041 | 0.009 | 0.010 | 0.029 | 0.004 | 0.023 | 0.041 | 0.013 | 0.019 | 0.031 | 0.013 | **0.021** |
| GeneCompass_C | 0.005 | 0.003 | 0.029 | 0.029 | 0.007 | 0.005 | 0.015 | 0.006 | 0.005 | 0.003 | 0.011 | 0.014 | 0.020 | 0.047 | **0.009** |
| RegFormer_C | 0.022 | 0.004 | 0.012 | 0.004 | 0.025 | 0.002 | 0.051 | 0.007 | 0.003 | 0.002 | 0.004 | 0.015 | 0.007 | 0.018 | **0.007** |
| scFoundation_C | 0.008 | 0.002 | 0.005 | 0.004 | 0.001 | 0.013 | 0.003 | 0.000 | 0.025 | 0.011 | 0.002 | 0.005 | 0.021 | 0.032 | **0.005** |
| Geneformer_A | 0.007 | 0.013 | 0.006 | 0.007 | 0.001 | 0.003 | 0.001 | 0.004 | 0.001 | 0.002 | 0.015 | 0.005 | 0.003 | 0.002 | **0.003** |
| CellOracle | 0.002 | 0.004 | 0.003 | 0.003 | 0.002 | 0.003 | 0.002 | 0.000 | 0.002 | 0.011 | 0.003 | 0.001 | 0.004 | 0.004 | **0.003** |
| scFoundation_S | 0.002 | 0.002 | 0.003 | 0.003 | 0.001 | 0.001 | 0.004 | 0.001 | 0.004 | 0.004 | 0.001 | 0.002 | 0.003 | 0.012 | **0.002** |
| scFoundation_A | 0.002 | 0.002 | 0.027 | 0.001 | 0.001 | 0.001 | 0.005 | 0.007 | 0.001 | 0.000 | 0.007 | 0.003 | 0.002 | 0.007 | **0.002** |
| GeneCompass_A | 0.002 | 0.005 | 0.002 | 0.002 | 0.001 | 0.002 | 0.008 | 0.001 | 0.003 | 0.001 | 0.001 | 0.002 | 0.002 | 0.002 | **0.002** |
| scGPT_A | 0.002 | 0.001 | 0.005 | 0.004 | 0.001 | 0.001 | 0.001 | 0.001 | 0.001 | 0.002 | 0.005 | 0.002 | 0.005 | 0.004 | **0.002** |
| scPRINT_A | 0.001 | 0.002 | 0.003 | 0.001 | 0.000 | 0.000 | 0.005 | 0.001 | 0.001 | 0.000 | 0.005 | 0.004 | 0.002 | 0.001 | **0.001** |

Table 9: Per-dataset perturbational Fisher odds ratio.

| Representation | Adam | Norm | K562 | RPE1 | Schm | SARS | MOLM | Endo | Ti-i | Ti-a | Drag | Leng | GWCD | WBrn | Median |
|---|---|---|---|---|---|---|---|---|---|---|---|---|---|---|---|
| CS-CORE | 2.79 | 2.47 | 1.09 | 2.32 | 1.98 | 9.44 | 1.63 | 7.80 | 22.16 | 4.90 | 0.57 | 0.43 | 0.61 | 46.94 | **2.39** |
| Geneformer_A | 2.96 | 4.90 | 1.19 | 1.17 | 3.27 | 5.24 | 2.84 | 4.37 | 9.41 | 3.65 | 1.91 | 7.91 | 2.42 | 4.20 | **3.46** |
| RegFormer_S | 1.66 | 2.30 | 1.34 | 1.64 | 2.18 | 2.39 | 0.64 | 2.87 | 26.03 | 2.35 | 2.73 | 3.38 | 1.65 | 1.89 | **2.24** |
| scFoundation_A | 1.80 | 2.58 | 3.10 | 2.64 | 2.00 | 2.64 | 1.22 | 6.11 | 6.73 | 3.12 | 1.04 | 1.94 | 1.91 | 1.61 | **2.29** |
| scPRINT_A | 2.96 | 2.72 | 1.44 | 1.29 | 1.38 | 2.81 | 2.34 | 1.08 | 1.87 | 4.71 | 2.26 | 2.84 | 1.50 | 1.51 | **2.07** |
| CellOracle | 1.74 | 3.17 | 0.99 | 1.00 | 3.29 | 7.20 | 2.82 | 1.50 | 1.75 | 3.07 | 1.44 | 3.24 | 1.72 | 12.60 | **2.29** |
| gene2vec | 2.45 | 1.57 | 0.97 | 2.52 | 2.37 | 3.39 | 1.10 | 2.21 | 7.13 | 3.09 | 0.80 | 2.37 | 1.10 | 0.43 | **2.29** |
| scGPT_S | 2.96 | 1.05 | 1.05 | 2.43 | 1.38 | 3.08 | 0.14 | 0.00 | 21.92 | 1.68 | 0.40 | 3.64 | 0.87 | 2.46 | **1.53** |
| scPRINT_C | 2.95 | 1.29 | 0.80 | 2.23 | 2.22 | 2.04 | 2.13 | 0.31 | 3.14 | 2.75 | 0.92 | 0.48 | 1.89 | 1.20 | **1.97** |
| GeneCompass_A | 3.52 | 2.79 | 0.97 | 1.14 | 2.97 | 2.28 | 2.09 | 6.01 | 7.06 | 4.80 | 1.95 | 2.19 | 1.85 | 1.08 | **2.23** |
| GenePT | 1.81 | 1.62 | 1.01 | 1.41 | 1.57 | 6.30 | 1.14 | 2.14 | 0.55 | 1.58 | 1.43 | 3.47 | 1.24 | 1.01 | **1.50** |
| scGPT_C | 3.95 | 1.84 | 1.19 | 1.94 | 1.87 | 2.53 | 2.03 | 11.20 | 2.41 | 1.62 | 1.54 | 3.66 | 0.97 | 2.80 | **1.99** |
| Geneformer_C | 1.57 | 1.15 | 1.00 | 1.39 | 1.07 | 2.35 | 1.33 | 0.65 | 7.64 | 4.59 | 0.50 | 1.35 | 1.58 | 0.82 | **1.34** |
| ESM | 1.26 | 1.16 | 0.69 | 1.40 | 1.48 | 1.63 | 1.63 | 0.05 | 5.92 | 2.05 | 0.95 | 1.93 | 1.28 | 4.04 | **1.44** |
| RegFormer_C | 2.50 | 1.03 | 0.71 | 1.25 | 1.55 | 1.91 | 1.93 | 0.58 | 2.81 | 3.10 | 1.48 | 1.71 | 1.36 | 1.28 | **1.51** |
| GeneCompass_C | 0.98 | 0.72 | 1.07 | 1.41 | 1.55 | 0.40 | 1.28 | 0.08 | 2.49 | 1.61 | 0.47 | 1.09 | 1.76 | 0.90 | **1.08** |
| scFoundation_S | 1.08 | 1.22 | 1.10 | 1.07 | 1.57 | 1.33 | 1.31 | 1.60 | 2.56 | 1.82 | 1.12 | 0.88 | 1.04 | 0.39 | **1.17** |
| Geneformer_S | 1.36 | 0.90 | 0.96 | 1.99 | 1.40 | 1.81 | 1.19 | 0.09 | 0.30 | 3.73 | 0.54 | 4.39 | 0.51 | 0.07 | **1.08** |
| scFoundation_C | 0.00 | 0.06 | 0.04 | 0.52 | 0.06 | 0.00 | 0.02 | 0.00 | 0.00 | 0.00 | 0.03 | 0.00 | 0.07 | 0.00 | **0.01** |
| GeneCompass_S | 0.94 | 0.80 | 0.81 | 1.76 | 1.06 | 2.90 | 0.86 | 0.00 | 0.00 | 2.29 | 0.36 | 1.68 | 0.59 | 0.00 | **0.83** |
| scGPT_A | 0.21 | 0.49 | 0.98 | 0.72 | 0.63 | 0.47 | 1.09 | 0.64 | 1.66 | 1.97 | 0.59 | 0.53 | 0.95 | 0.62 | **0.64** |

Table 10: Per-dataset perturbational AUROC.

| Representation | Adam | Norm | K562 | RPE1 | Schm | SARS | MOLM | Endo | Ti-i | Ti-a | Drag | Leng | GWCD | WBrn | Median |
|---|---|---|---|---|---|---|---|---|---|---|---|---|---|---|---|
| CS-CORE | 0.567 | 0.434 | 0.465 | 0.495 | 0.513 | 0.510 | 0.512 | 0.297 | 0.731 | 0.539 | 0.402 | 0.498 | 0.456 | 0.719 | **0.504** |
| Geneformer_A | 0.477 | 0.494 | 0.468 | 0.493 | 0.515 | 0.440 | 0.517 | 0.394 | 0.425 | 0.449 | 0.520 | 0.676 | 0.509 | 0.420 | **0.485** |
| RegFormer_S | 0.504 | 0.526 | 0.580 | 0.512 | 0.539 | 0.521 | 0.262 | 0.624 | 0.750 | 0.515 | 0.521 | 0.544 | 0.520 | 0.541 | **0.523** |
| scFoundation_A | 0.507 | 0.418 | 0.606 | 0.625 | 0.494 | 0.514 | 0.423 | 0.520 | 0.263 | 0.422 | 0.424 | 0.330 | 0.507 | 0.630 | **0.501** |
| scPRINT_A | 0.514 | 0.507 | 0.501 | 0.500 | 0.430 | 0.449 | 0.547 | 0.521 | 0.444 | 0.455 | 0.506 | 0.539 | 0.517 | 0.278 | **0.504** |
| CellOracle | 0.541 | 0.678 | 0.487 | 0.557 | 0.644 | 0.780 | 0.600 | 0.398 | 0.623 | 0.620 | 0.569 | 0.609 | 0.644 | 0.714 | **0.615** |
| gene2vec | 0.510 | 0.520 | 0.497 | 0.535 | 0.522 | 0.494 | 0.416 | 0.279 | 0.489 | 0.535 | 0.490 | 0.529 | 0.464 | 0.426 | **0.495** |
| scGPT_S | 0.600 | 0.566 | 0.511 | 0.543 | 0.539 | 0.660 | 0.426 | 0.500 | 0.535 | 0.481 | 0.197 | 0.580 | 0.447 | 0.481 | **0.523** |
| scPRINT_C | 0.550 | 0.508 | 0.430 | 0.526 | 0.514 | 0.512 | 0.483 | 0.239 | 0.342 | 0.431 | 0.471 | 0.481 | 0.435 | 0.606 | **0.482** |
| GeneCompass_A | 0.621 | 0.475 | 0.361 | 0.466 | 0.503 | 0.630 | 0.362 | 0.505 | 0.338 | 0.500 | 0.540 | 0.480 | 0.424 | 0.391 | **0.477** |
| GenePT | 0.560 | 0.502 | 0.516 | 0.529 | 0.525 | 0.626 | 0.424 | 0.533 | 0.435 | 0.536 | 0.505 | 0.634 | 0.495 | 0.441 | **0.520** |
| scGPT_C | 0.532 | 0.472 | 0.565 | 0.521 | 0.524 | 0.489 | 0.470 | 0.274 | 0.300 | 0.421 | 0.497 | 0.462 | 0.436 | 0.400 | **0.471** |
| Geneformer_C | 0.494 | 0.484 | 0.517 | 0.494 | 0.460 | 0.514 | 0.539 | 0.515 | 0.448 | 0.520 | 0.487 | 0.680 | 0.492 | 0.754 | **0.504** |
| ESM | 0.513 | 0.533 | 0.512 | 0.540 | 0.495 | 0.502 | 0.461 | 0.425 | 0.561 | 0.534 | 0.456 | 0.715 | 0.490 | 0.589 | **0.512** |
| RegFormer_C | 0.546 | 0.512 | 0.507 | 0.517 | 0.453 | 0.525 | 0.450 | 0.608 | 0.423 | 0.462 | 0.461 | 0.341 | 0.492 | 0.472 | **0.482** |
| GeneCompass_C | 0.470 | 0.470 | 0.537 | 0.526 | 0.475 | 0.568 | 0.483 | 0.411 | 0.449 | 0.459 | 0.420 | 0.619 | 0.494 | 0.445 | **0.472** |
| scFoundation_S | 0.469 | 0.520 | 0.494 | 0.506 | 0.492 | 0.465 | 0.502 | 0.622 | 0.432 | 0.484 | 0.483 | 0.381 | 0.501 | 0.447 | **0.488** |
| Geneformer_S | 0.436 | 0.499 | 0.455 | 0.513 | 0.488 | 0.533 | 0.598 | 0.420 | 0.499 | 0.519 | 0.483 | 0.561 | 0.459 | 0.355 | **0.493** |
| scFoundation_C | 0.500 | 0.365 | 0.283 | 0.431 | 0.253 | 0.500 | 0.554 | 0.500 | 0.500 | 0.500 | 0.441 | 0.500 | 0.481 | 0.500 | **0.500** |
| GeneCompass_S | 0.476 | 0.481 | 0.447 | 0.497 | 0.470 | 0.655 | 0.368 | 0.500 | 0.500 | 0.447 | 0.330 | 0.757 | 0.446 | 0.500 | **0.479** |
| scGPT_A | 0.417 | 0.463 | 0.487 | 0.486 | 0.423 | 0.250 | 0.491 | 0.739 | 0.385 | 0.473 | 0.412 | 0.377 | 0.462 | 0.446 | **0.454** |

Table 11: Per-dataset regulator-level enrichment rate.

| Representation | Adam | Norm | K562 | RPE1 | Schm | SARS | MOLM | Endo | Ti-i | Ti-a | Drag | Leng | GWCD | WBrn | Median |
|---|---|---|---|---|---|---|---|---|---|---|---|---|---|---|---|
| CS-CORE | 0.162 | 0.151 | 0.085 | 0.404 | 0.134 | 0.102 | 0.037 | 0.010 | 0.009 | 0.069 | 0.012 | 0.000 | 0.025 | 0.026 | **0.053** |
| Geneformer_A | 0.411 | 0.552 | 0.016 | 0.009 | 0.435 | 0.036 | 0.136 | 0.011 | 0.008 | 0.119 | 0.010 | 0.077 | 0.086 | 0.002 | **0.057** |
| RegFormer_S | 0.100 | 0.091 | 0.006 | 0.085 | 0.180 | 0.011 | 0.000 | 0.010 | 0.014 | 0.026 | 0.009 | 0.026 | 0.014 | 0.000 | **0.014** |
| scFoundation_A | 0.210 | 0.229 | 0.010 | 0.090 | 0.297 | 0.014 | 0.091 | 0.016 | 0.004 | 0.091 | 0.019 | 0.038 | 0.031 | 0.000 | **0.035** |
| scPRINT_A | 0.599 | 0.444 | 0.003 | 0.007 | 0.168 | 0.031 | 0.303 | 0.005 | 0.000 | 0.169 | 0.046 | 0.026 | 0.041 | 0.000 | **0.036** |
| CellOracle | 0.012 | 0.226 | 0.000 | 0.007 | 0.122 | 0.021 | 0.083 | 0.000 | 0.000 | 0.000 | 0.000 | 0.000 | 0.006 | 0.000 | **0.003** |
| gene2vec | 0.170 | 0.047 | 0.000 | 0.217 | 0.167 | 0.019 | 0.000 | 0.005 | 0.004 | 0.025 | 0.000 | 0.012 | 0.018 | 0.000 | **0.015** |
| scGPT_S | 0.262 | 0.056 | 0.064 | 0.278 | 0.072 | 0.009 | 0.000 | 0.000 | 0.004 | 0.000 | 0.000 | 0.013 | 0.017 | 0.000 | **0.011** |
| scPRINT_C | 0.414 | 0.027 | 0.012 | 0.239 | 0.226 | 0.004 | 0.070 | 0.000 | 0.002 | 0.045 | 0.010 | 0.000 | 0.092 | 0.000 | **0.020** |
| GeneCompass_A | 0.260 | 0.396 | 0.000 | 0.002 | 0.296 | 0.008 | 0.258 | 0.005 | 0.008 | 0.129 | 0.009 | 0.026 | 0.059 | 0.000 | **0.017** |
| GenePT | 0.077 | 0.030 | 0.023 | 0.052 | 0.032 | 0.040 | 0.000 | 0.000 | 0.000 | 0.000 | 0.000 | 0.000 | 0.002 | 0.000 | **0.001** |
| scGPT_C | 0.635 | 0.211 | 0.066 | 0.237 | 0.227 | 0.025 | 0.161 | 0.015 | 0.000 | 0.023 | 0.009 | 0.013 | 0.042 | 0.000 | **0.033** |
| Geneformer_C | 0.040 | 0.003 | 0.018 | 0.116 | 0.043 | 0.016 | 0.000 | 0.000 | 0.004 | 0.080 | 0.000 | 0.000 | 0.016 | 0.002 | **0.010** |
| ESM | 0.011 | 0.003 | 0.000 | 0.030 | 0.000 | 0.010 | 0.042 | 0.000 | 0.000 | 0.004 | 0.000 | 0.000 | 0.001 | 0.000 | **0.000** |
| RegFormer_C | 0.520 | 0.000 | 0.000 | 0.061 | 0.101 | 0.010 | 0.130 | 0.000 | 0.006 | 0.085 | 0.010 | 0.000 | 0.014 | 0.000 | **0.010** |
| GeneCompass_C | 0.021 | 0.000 | 0.023 | 0.082 | 0.065 | 0.000 | 0.014 | 0.000 | 0.000 | 0.027 | 0.000 | 0.012 | 0.028 | 0.003 | **0.013** |
| scFoundation_S | 0.000 | 0.000 | 0.000 | 0.000 | 0.005 | 0.000 | 0.000 | 0.000 | 0.000 | 0.004 | 0.000 | 0.000 | 0.000 | 0.000 | **0.000** |
| Geneformer_S | 0.045 | 0.000 | 0.032 | 0.154 | 0.055 | 0.002 | 0.000 | 0.000 | 0.000 | 0.000 | 0.000 | 0.000 | 0.003 | 0.000 | **0.000** |
| scFoundation_C | 0.000 | 0.000 | 0.016 | 0.034 | 0.000 | 0.000 | 0.000 | 0.000 | 0.000 | 0.000 | 0.000 | 0.000 | 0.000 | 0.000 | **0.000** |
| GeneCompass_S | 0.004 | 0.007 | 0.021 | 0.138 | 0.029 | 0.000 | 0.000 | 0.000 | 0.000 | 0.000 | 0.000 | 0.000 | 0.003 | 0.000 | **0.000** |
| scGPT_A | 0.000 | 0.007 | 0.000 | 0.000 | 0.000 | 0.000 | 0.000 | 0.000 | 0.000 | 0.011 | 0.009 | 0.000 | 0.003 | 0.000 | **0.000** |

### Appendix J Robustness analyses

#### J.1 Sensitivity to Benchmark Settings

We evaluated the sensitivity of representation rankings to the benchmark choices: network size, GO similarity threshold, the number of cells and the cell population used for contextual extraction, and similarity definition. Rankings were recomputed under each setting and compared using Spearman correlation. Table 12 summarizes the results, and Figure 6 visualizes the first four analyses. Each axis varies one choice from the reported configuration (10,000 edges, Jaccard $> 0.3$). CellOracle is excluded because its networks do not reach 10,000 or 20,000 edges on most datasets.

Table 12: Stability of representation rankings across benchmark settings. Agreement is the mean pairwise Spearman correlation between rankings; rank span summarizes movement across settings.

| Axis | Settings | Representations | Agreement | Rank span (med / max) |
|---|---|---|---|---|
| Network size | 5,000 / 10,000 / 20,000 edges | 20 | 0.943 | 1.0 / 6 |
| GO reference | Jaccard $> 0.1$ / $> 0.3$ | 20 | 0.910 | 1.0 / 6 |
| Cells subsampled | 250 / 500 / 1,000 / 2,000 / 4,000 | 11 | 0.902 | 2.0 / 4 |
| Cell population | mixed 1,000 / control 1,000 / all control | 11 | 0.876 | 1.5 / 4 |
| Similarity | cosine / −PC1 / gaussianized | 8 | 0.873 | 1.0 / 3 |

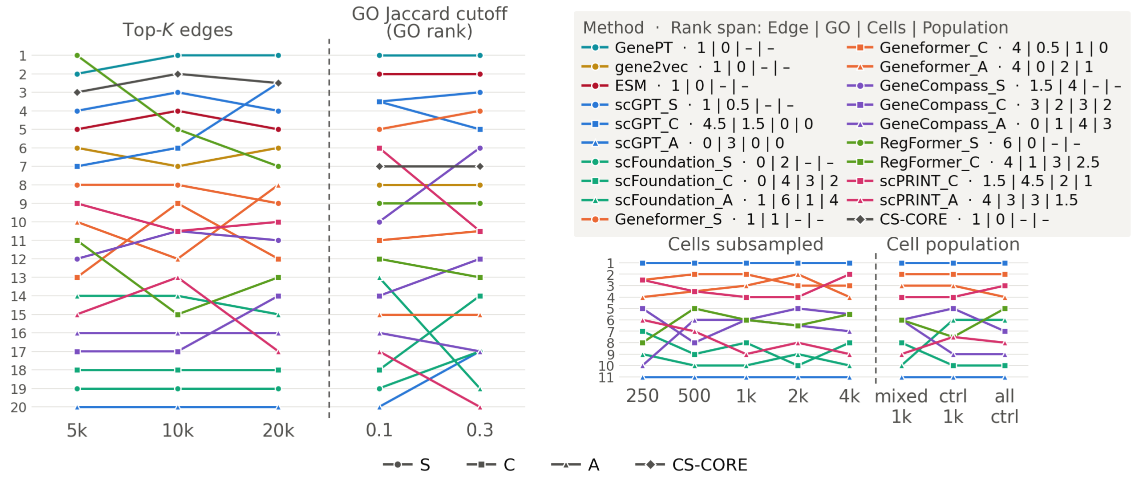


Figure 6: Rank stability across network size, GO Jaccard threshold, contextual cell-subsample size, and cell population. Lower ranks indicate better performance; the GO-threshold panel uses GO-specific ranks, and the cell-subsampling and cell-population analyses include only contextual and attention-based representations.

**Network size.** Rankings were stable across networks with 5,000, 10,000, and 20,000 edges (mean pairwise Spearman $\rho = 0.943$). The median rank span was 1.0 positions, suggesting that broad trends are stable while small rank differences should not be over-interpreted.

**GO similarity threshold.** Using GO Jaccard thresholds of 0.1 and 0.3 produced similar GO rankings ($\rho = 0.910$), despite the stricter reference at 0.3. Both thresholds are therefore reported as a robustness check.

**Cell-subsample size.** Contextual and attention representations were recomputed from nested random subsamples of 250, 500, 1,000, 2,000, and 4,000 cells drawn from all cells of each dataset. Rankings remained stable across this range (mean pairwise $\rho = 0.902$), with no representation moving by more than four positions, indicating that rankings of the contextual sources are insensitive to the number of cells used for extraction.

**Cell population.** The main analysis extracts contextual representations and attention scores from control cells only. To test whether including perturbed cells changes the comparison, we also extracted them from 1,000 cells sampled from all cells of each dataset, most of them perturbed, and from 1,000 control cells (all control cells where fewer are available). Rankings of the 11 contextual sources agreed across the three inputs (mean pairwise $\rho = 0.876$), with no source moving by more than four positions.

**Similarity definition.** For vector representations, we compared plain cosine similarity with cosine after removing the leading principal component and with an additional rank-inverse-normal transformation. Cosine and PC1-removed cosine gave identical rankings, and the gaussianized variant moved no representation by more than three positions (mean pairwise $\rho = 0.873$; Geneformer's static embedding rose from sixth to third), so the main benchmark reports the unmodified cosine similarity.

Overall, the broad representation ordering and group-level conclusions are stable across these design choices, whereas differences of only a few rank positions should not be interpreted as robust differences between individual representations.

#### J.2 Sensitivity to Dataset Characteristics

We next tested whether broad biological characteristics of the Perturb-seq datasets systematically influenced benchmark scores. For each dataset and metric, scores were first averaged across the three network sizes (5k, 10k, and 20k edges) within each representation and then across the 20 representations, yielding one dataset-level value per metric. Datasets were stratified by tissue (brain, $n = 5$; blood, $n = 6$), cell system (cell line, $n = 6$; primary/iPSC, $n = 8$), and perturbation type (CRISPRi, $n = 9$; CRISPRa, $n = 3$). Group differences were assessed using two-sided Mann–Whitney $U$ tests, with Cliff's $\delta$ as the effect size.

None of the comparisons remained significant after Benjamini–Hochberg correction (Table 13). Two comparisons showed nominal differences: regulator-level enrichment between brain and blood datasets ($\delta = -0.87$, $p = 0.017$) and GO AUPRC between CRISPRi and CRISPRa datasets ($\delta = +0.85$, $p = 0.036$), but neither survived multiple-testing correction. Overall, we found no statistically significant evidence that the benchmark metrics were systematically driven by the tested tissue, cell-system, or perturbation-type differences.

Table 13: Association between dataset-level biological characteristics and benchmark metrics. Effect sizes are reported as Cliff's $\delta = P(X > Y) - P(X < Y)$, where $X$ and $Y$ denote the first and second groups listed, respectively. Bold values indicate nominal $p < 0.05$; no comparison remained significant after Benjamini–Hochberg correction.

| **Metric** | **Tissue** | | **Cell system** | | **Perturbation** | |
|---|---|---|---|---|---|---|
| | brain ($n = 5$) vs. blood ($n = 6$) | | cell line ($n = 6$) vs. primary/iPSC ($n = 8$) | | CRISPRi ($n = 9$) vs. CRISPRa ($n = 3$) | |
| | $\delta$ | $p$ | $\delta$ | $p$ | $\delta$ | $p$ |
| *GO, Jaccard* $> 0.3$ | | | | | | |
| Fisher OR | −0.13 | 0.792 | −0.08 | 0.852 | +0.63 | 0.145 |
| AUROC | −0.47 | 0.247 | +0.54 | 0.108 | +0.41 | 0.373 |
| AUPRC | +0.40 | 0.329 | +0.04 | 0.950 | +0.85 | **0.036** |
| *Perturb-seq* | | | | | | |
| DE Fisher OR | +0.67 | 0.082 | −0.42 | 0.228 | −0.19 | 0.727 |
| DE AUROC | −0.07 | 0.931 | +0.46 | 0.181 | +0.04 | 1.000 |
| Regulator-level enrichment | −0.87 | **0.017** | +0.62 | 0.059 | −0.70 | 0.100 |

### Appendix K Computational cost

All runtimes are wall-clock measurements from the benchmark cluster. Contextual representation extraction is the only GPU stage and uses one NVIDIA A30 (24 GB) with 8 CPU cores and 64 GB host memory. Differential-expression testing, CS-CORE, CellOracle, network construction, and evaluation are CPU-only and use 8 cores with up to 128 GB memory. No stage requires multiple GPUs or distributed execution.

Static representations are read directly from released embedding files or checkpoint token tables and therefore incur negligible inference cost. For contextual representations, we measured runtime across six foundation models using 250-4,000 subsampled cells and approximately 1,000 genes (Table 14).

Table 14: Runtime of contextual representation extraction on one NVIDIA A30.

| | **Checkpoint** | **Load** | **Seconds per dataset** | | | | | **Slope** |
|---|---|---|---|---|---|---|---|---|
| **Model** | (MB) | (s) | **250** | **500** | **1,000** | **2,000** | **4,000** | |
| scPRINT | 211 | 30 | 3.3 | 3.7 | 2.7 | 6.6 | 10.8 | 0.43 |
| GeneCompass | 1099 | $< 5$ | 4.5 | 4.8 | 8.8 | 17.0 | 33.2 | 0.76 |
| scGPT | 196 | 127 | 11.7 | 14.4 | 14.4 | 33.9 | 59.8 | 0.59 |
| Geneformer | 398 | $< 5$ | 12.9 | 14.6 | 20.0 | 26.9 | 42.9 | 0.43 |
| RegFormer | 189 | 6 | 7.0 | 12.1 | 24.1 | 47.3 | 93.5 | 0.94 |
| scFoundation | 1366 | $< 5$ | 43.6 | 85.9 | 170.0 | 342.2 | 680.0 | 0.99 |
| **All six, 14 datasets** | | | **24 min** | **38 min** | **68 min** | **2.2 h** | **4.3 h** | |

Runtime increases with cell count and varies substantially across models (Figure 7). At 4,000 cells, extraction ranges from 10.8 s for scPRINT to 680 s for scFoundation, while all models complete within one hour per dataset. In the main benchmark, extraction uses all post-QC control cells; running all six contextual models requires approximately 5.6 GPU-hours, 3.9 of them for scFoundation. The full cell-subsampling sweep requires about 8.6 GPU-hours.

The SCEPTRE differential-expression reference requires approximately 17 CPU-hours in total

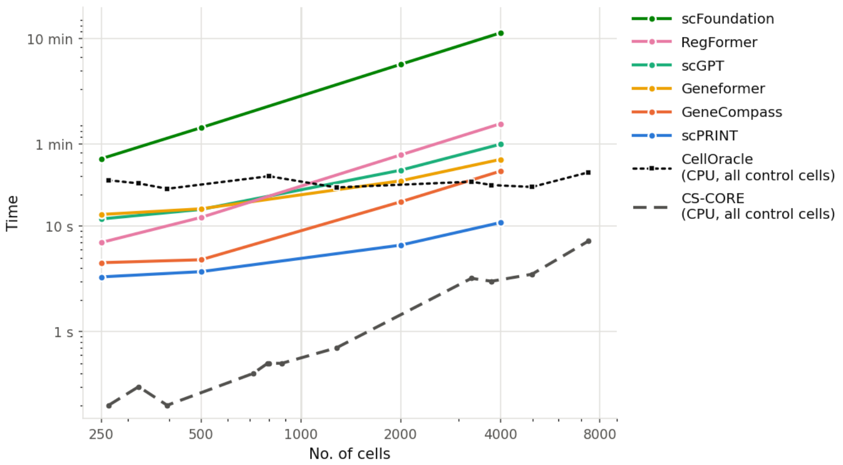


Figure 7: Runtime scaling of contextual foundation models and CS-CORE. Solid lines show mean runtime for six contextual models on one NVIDIA A30; the dashed lines show CS-CORE and CellOracle using eight CPU cores and all available control cells. Both axes are logarithmic.

but is computed once per dataset and reused by all methods. CS-CORE is substantially cheaper, requiring 0.2-234 s per dataset on eight CPU cores. CellOracle's runtime remains stable at approximately 22-32 s below 7,000 cells, dominated by fixed computational costs such as loading TF-target priors. Beyond 10,000 cells, runtime increases as ridge regression and KNN imputation become more computationally expensive. Network construction and evaluation are also CPU-only, requiring approximately 0.6 minutes per representation per dataset across the three network sizes in the reported cosine arm.

Overall, the main benchmark requires approximately 5.6 GPU-hours for contextual representation extraction, while static representations require no GPU inference. The remaining computational cost is dominated by CPU-based differential-expression testing and network evaluation.
